# A dual genomic-epigenomic map of clonal evolution in grapevine

**DOI:** 10.64898/2026.01.09.698625

**Authors:** Paolo Callipo, Hannah Robinson, Maximilian Schmidt, Kai P. Voss-Fels

## Abstract

Grapevine is one of the world’s oldest and most economically important perennial crops and has been vegetatively propagated for centuries to millennia. This long history of clonal propagation has generated substantial intra-varietal diversity, but its molecular basis has remained difficult to resolve with short-read sequencing, which misses large structural variation and DNA base modifications. Here, we generated a high-quality, phased diploid reference genome for the elite cultivar Pinot noir and integrate it with Oxford Nanopore sequencing of 23 distinct clones to build an integrated, genome-wide map of clonal genetic and epigenetic variation in *Vitis vinifera*. The genome assembly reveals a deep history of ancient inbreeding, with approximately 12% of the genome occurring in extended runs of homozygosity. Across molecular layers, we observe contrasting evolutionary dynamics. Somatic genetic variation (67,277 SNPs and 4,037 SVs) is dominated by rare variants and strongly depleted from coding regions, consistent with strong purifying selection. A substantial fraction of the structural variation among clones is mechanistically linked to instability in repetitive DNA, particularly centromeric repeats. In contrast, CG context epigenetic variation is abundant (250,382 differentially methylated cytosines) and strongly enriched within gene bodies, highlighting it as a major source of clonal divergence. Most importantly, CG methylation patterns alone reconstruct clonal phylogenetic relationships with high fidelity, closely mirroring the SNP-based lineages. This congruence, absent in transient non-CG methylation, indicates that stable CG epialleles are mitotically inherited and preserve a persistent record. Together, our results show that clonal identity in grapevine is jointly encoded by genome and methylome, with stable CG methylation capturing propagation history at near-genetic resolution.

## Introduction

Grapevine (*Vitis vinifera* L.) is one of the earliest crops domesticated during the Neolithic revolution, establishing it as a mainstay in global horticulture (McGovern 2003; Dong et al. 2023). Spanning millennia of cultivation, it is now one of the world’s most economically important fruit crops, generating a global farm gate value estimated at about 68 billion dollars annually (Alston and Sambucci 2019). Viticulture, driven by the production of high-value commodities such as wine, table grapes, and raisins, is a critical component of agricultural economies across nearly every continent (OIV 2025).

This long history of cultivation, coupled with its perennial and clonal lifecycle, has positioned grapevine as an exceptional model for studying evolutionary genetics, from the initial domestication of key fruit traits (This et al. 2006; Zhou et al. 2017), to the long-term accumulation of somatic mutations and epigenetic modifications within ancient clonal lineages (Vondras et al. 2019, Varela et al. 2021, Xiao et al. 2025, Callipo et al. 2025).

Despite its relatively compact genome of approximately 500 Mb, the genomic architecture of grapevine is exceptionally complex (Jaillon et al. 2007; Velasco et al. 2007). This complexity is a direct legacy of its outcrossing nature coupled with vegetative propagation practices to maintain highly heterozygous, elite genotypes (Zhou et al. 2017). Recent analyses confirm high heterozygosity (Zhou et al. 2019), extreme hemizygosity, with up to 25% of content exclusive to one haplotype (Peng et al. 2025), and in some old cultivars, extensive Runs of Homozygosity (RoH), derived from ancient inbreeding (Roach et al. 2018). This interspersed landscape of heterozygosity, hemizygosity, and homozygosity represents a fundamental challenge to traditional genomic studies. Furthermore, the role of DNA methylation within these structurally complex regions remains largely unexplored, with only one recent study beginning to address the distinct epigenetic patterns of hemizygous genes (Peng et al. 2025).

The practice of clonal propagation carried out for hundreds to thousands of years, essential for preserving the genetic and phenotypic integrity of ancient cultivars (Pelsy 2010), paradoxically functions as an engine for their diversification. Over millennia, the accumulation of somatic mutations and epigenetic modifications creates a vast array of molecularly distinct individuals, or clones, from a single founder (Callipo et al. 2025). Pinot noir serves as the ideal model to study this process. It is both a foundational parent for many modern cultivars like Chardonnay, Auxerois, and Gamay (Regner et al. 2000, Lacombe et al. 2013) and remains one of the world’s most important cultivars for wine production. Its extensive propagation history, spanning nearly a millennium (Ramos-Madrigal et al. 2019), has generated substantial somatic variation, including famous color mutants like Pinot gris and Pinot blanc (Vezzulli et al. 2012). Crucially, this long-term somatic evolution has produced valuable intra-varietal variation for key traits, providing an invaluable resource for both applied breeding and scientific discovery (Robinson et al. 2025).

Even though viticulture and grapevine breeding has systematically exploited intra-varietal variation to release improved varietal clones (van Houten et al. 2020, Portu et al. 2024, Carvalho et al. 2025), a comprehensive understanding of the molecular factors that drive clonal diversity has remained elusive. In recent years, whole-genome re-sequencing of clonal populations in key cultivars, including Pinot noir, Chardonnay, Nebbiolo, and Zinfandel, has begun to unravel the molecular basis of this diversity, primarily by identifying clone-specific single nucleotide polymorphisms (SNPs) and small InDels, which are found primarily in intergenic regions and are often predicted to have a low functional impact (Gambino et al. 2017; Roach et al. 2018; Vondras et al. 2019; Urra et al. 2023; Xiao et al. 2025). Nevertheless, these pioneering genomic efforts have been largely constrained by the technical limitations of the short-read sequencing technologies they employed. These methods are often unable to resolve larger, more complex genomic rearrangements, and consequently, the functional impact of structural variants (SVs) remains poorly understood. Additionally, the widespread use of a haploid reference genome introduces analytical biases, such as the overestimation of heterozygous loci or the failure to observe true somatic variants (Wang et al. 2024).

Furthermore, an entire layer of molecular variation, the epigenetic landscape of DNA methylation, has remained almost entirely unexplored, mainly due to technical limitations. While a few studies have implicated the importance of DNA methylation in clonal diversification (Ocaña et al. 2013) and environmental plasticity (Varela et al. 2021), a comprehensive, genome-wide view of its role in shaping clonal phenotypes represents a major, largely unexplored frontier in grapevine genetics, and more broadly other clonally propagated crops.

Recent advances in long-read sequencing, particularly Oxford Nanopore Technology (ONT), provide a unified solution to these multifaceted challenges. The capacity of long reads to span complex repetitive and heterozygous regions enables the direct phasing of genomic reads and so the accurate characterization of somatic variants, including large structural variants, that were previously unresolved by short-read methods. Furthermore, by analysing native DNA,

ONT concurrently provides a base-resolution view of the methylome. As such, ONT provides a powerful, integrated methodology that has the potential to dissect the complex molecular architecture of intra-varietal variation in grapevine (Schmidt et al. 2025).

Here, we address these gaps by generating a new phased diploid genome assembly and its methylome for the cultivar Pinot noir using PacBio HiFi and Oxford Nanopore Ultra-long read sequencing. Leveraging this high-resolution genome, we performed the first integrated analysis of both genomic and epigenomic variation across a panel of 23 Pinot clones using Nanopore sequencing. Our analysis highlights several key features: First, we uncover a deep history of ancient inbreeding, with approximately 12% of the Pinot noir genome occurring in a state of complete homozygosity. Second, we show that large structural variation between clones is non-randomly distributed, with repetitive DNA acting as preferential sites for somatic rearrangement. Third, we find that CG-DMCs are significantly enriched in coding regions, indicating that stable epigenetic variation represents an important component of clonal diversification. Finally, we show that the high-fidelity phylogenetic relationships inferred from SNPs are closely mirrored by patterns of mitotically stable CG-DMCs. This work supports a model in which clonal identity in perennial crops reflects the combined contributions of genome and epigenome.

## Results

### Establishing a Diploid, Chromosome-Scale Reference Genome to Decode Clonal Evolution in Pinot noir

To provide a precise reference for our Clonal Pinot noir population, we generated a de novo assembly of the clone ‘20-13 Gm’ using a hybrid approach combining PacBio HiFi and Oxford Nanopore ultra-long (>50kb) reads (Table S1). We successfully resolved the genome into two fully phased, chromosome-scale pseudo-haplotypes (PN_1 and PN_2), each spanning approximately 495 Mb. Both haplotypes demonstrate exceptional contiguity, achieving near chromosome-scale contig N50s of 26.3 Mb. High completeness is confirmed by both gene content, with a BUSCO score of over 98% for both haplotypes (Figure S1), and by k-mer representation, at 99.5% for the combined diploid assembly. The base-level accuracy of the assembly is exceedingly high, with a Merqury-derived Quality Value (QV) for both Haplotypes of 75, corresponding to an estimated error rate of less than one in 30 million bases. The overall structural integrity was further assessed with the Clipping Reveals Assembly Quality (CRAQ) tool (Li et al. 2023), yielding an Overall Assembly Quality Index (S-AQI) of 98 and a Regional Assembly Quality Index (R-AQI) of 99, respectively, indicating a near-complete, phased and correctly assembled genome structure (Table S2). We annotated approximately 35,000 high-confidence protein-coding genes per haplotype (Figure 1, Table S3), by combining ab initio predictions with high-quality gene models from PN40024.v4 (Velt et al. 2023). Transposable elements (TEs) comprise approximately 44% of the genome, consisting of more than 300,000 individual elements per haplotype (Figure 1, Table S4). The evolutionary history of TEs is characterized by a massive, recent amplification burst of LTR/Copia and LTR/Gypsy retrotransposons, indicated by a large proportion of elements with low sequence divergence (<5%) (Figure S2). This recent activity contrasts with a more ancient and diverse population of older TEs. The near-identical divergence profiles of the two haplotypes confirm their shared evolutionary history and validate the quality of the phasing (Figure S2). Leveraging the native Oxford Nanopore signal, we also generated base-resolution methylation profiles for each haplotype. Methylation levels were calculated, among the two haplotypes, for: 15 million sites in the CG context, 25 million sites in the CHG context, and 277 million in the CHH context. Spatially, the distribution of this methylation followed a canonical plant pattern, with hypermethylated regions correlating strongly with TE-dense areas and pericentromeric regions (Figure 1). This integrated resource provides a robust foundation for dissecting clonal variation.

**Figure 1.**
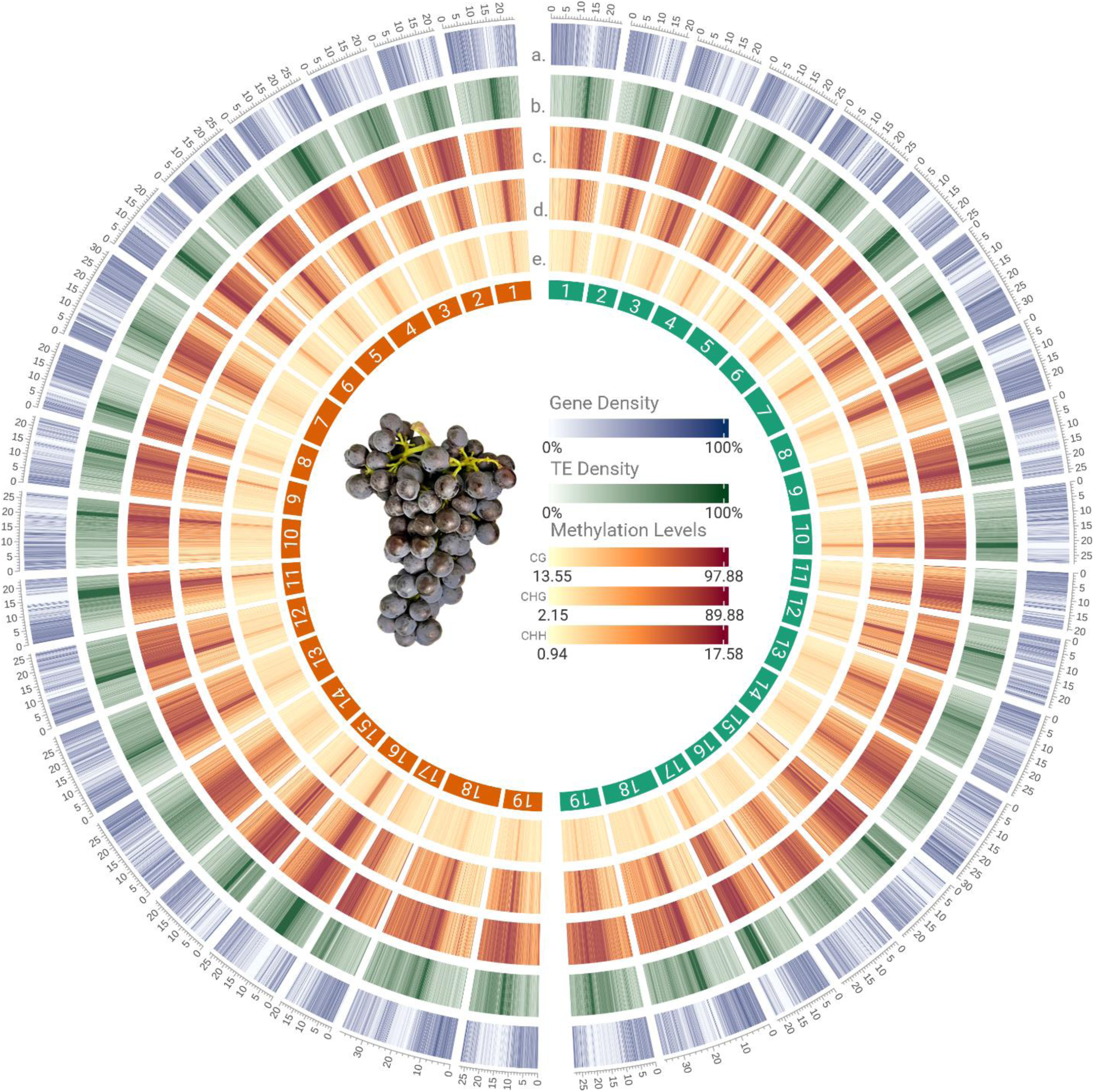
Genomic and Epigenomic Landscape of the Diploid Pinot noir Genome. Integrated visualization of the 19 diploid chromosome pairs with Haplotype 1 (labels in green, e.g., PN1_1) and Haplotype 2 (labels in orange, e.g., PN1_2). The outermost track indicates genomic scale (Mb). Inner concentric tracks display features calculated in 100kb non-overlapping bins: **(a)** Gene density; **(b)** Transposable element (TE) density; and DNA methylation levels in the **(c)** CG, **(d)** CHG, and **(e)** CHH contexts.

### Large-scale hemizygosity and extended runs of homozygosity reveal a structurally asymmetric Pinot noir genome

Whole-genome alignment and variant calling of the two pseudo-haplotypes using SyRI (Goel et al. 2019) revealed a dense landscape of variation, comprising 3.4 million single nucleotide polymorphisms (SNPs), over 500,000 InDels, and nearly 1,500 large structural variants, including 66 major inversions (Figure 2, Table S5). A prominent feature of the diploid genome, visible as distinct black zones in the SNP density track, is the presence of extensive runs of homozygosity (RoH) (Figure 2). These homozygous tracts collectively cover approximately 12% of the genome, totalling ∼60 Mb. While these RoH are distributed across most chromosomes, often concentrating in sub telomeric regions, several are exceptionally large. Notably, chromosome 14 contains a single homozygous block of 13.3 Mb extending from the telomere towards the centromere, and a 10 Mb RoH is present in the pericentromeric region of chromosome 5 (Figure 2). Beyond these linear variants, our analysis partitioned the diploid genome into its three primary structural states (Table S6). While the majority of the genome is heterozygous (62%), the remaining 26% is classified as functionally hemizygous, a classification which include regions that lack a direct syntenic or allelic counterpart in the opposing haplotype (see detailed definition in the Supplementary Information). These hemizygous portions are overwhelmingly repetitive, with transposable elements comprising 84% of their length and a corresponding gene content of just 13%. In contrast, heterozygous and homozygous regions contain 42% and 37% gene content respectively. This demonstrates that hemizygous regions constitute a genomic compartment characterized by transposable element accumulation and low gene density. Genes located in the heterozygous and homozygous regions are structurally conserved, typically having a syntenic counterpart on the opposite haplotype and few duplicated copies. In both cases, 98% of genes have a direct syntenic counterpart, and only 24% have duplicated copies within the same haplotype. In stark contrast, genes within hemizygous regions are defined by high rates of turnover. Here, 81.8% of genes have at least one duplicated copy on the same haplotype (Figure S4). Furthermore, genes within hemizygous regions display a trend towards shorter lengths and a greater proportion of single-exon models when compared to those in the core genomic partitions (Figure S5). The sequence divergence profiles of transposable elements also differed between the partitions. The distribution for TEs in hemizygous regions displayed a bimodal pattern, with a sharp peak corresponding to recent insertions (<5% divergence). In contrast, the profiles for TEs in the heterozygous and homozygous regions were largely unimodal and lacked this prominent peak of young elements (Figure S6).

**Figure 2.**
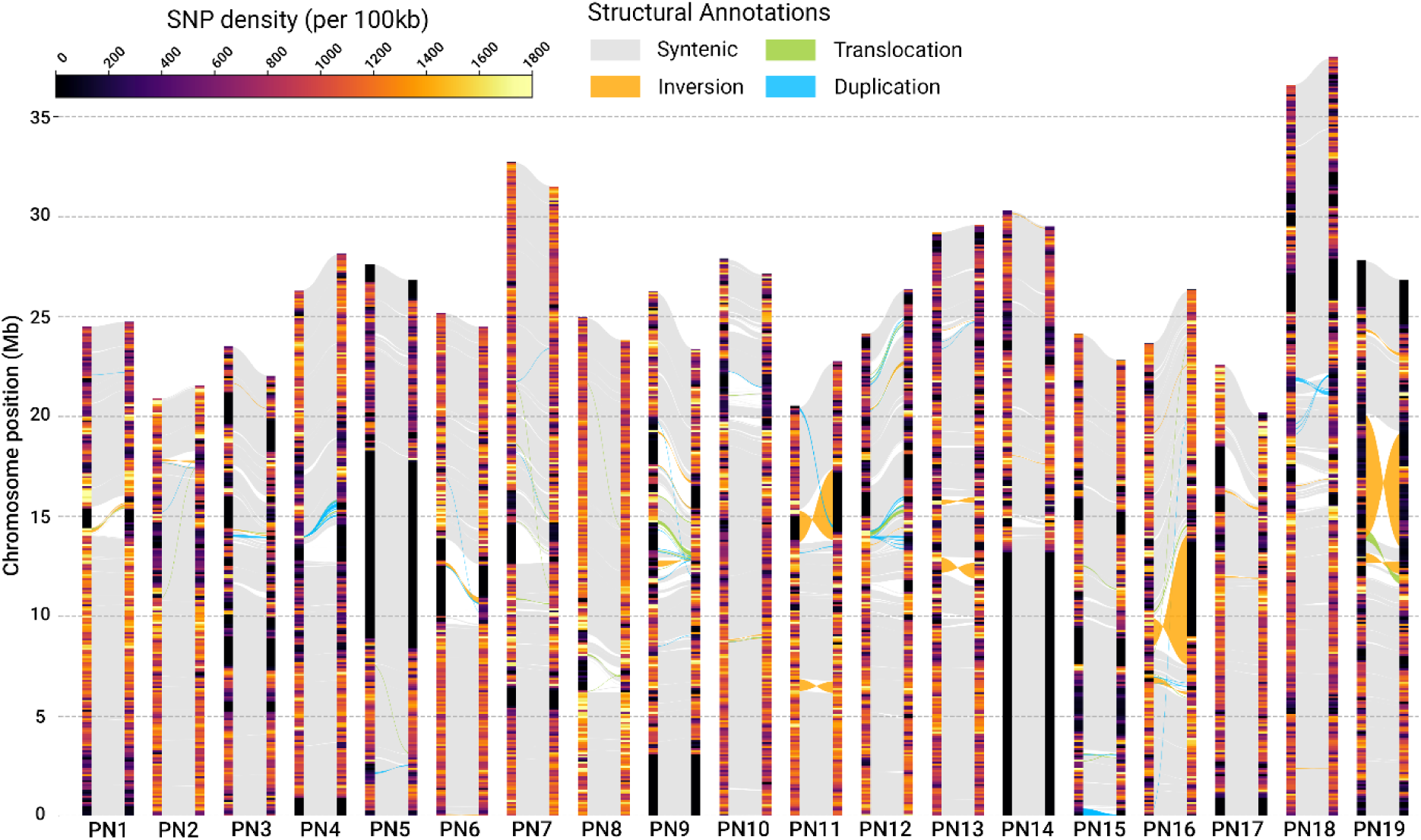
Genome-wide comparison of haplotype structure and SNP density. Comparative analysis of the two haplotypes across 19 chromosomes (PN1-PN19), with chromosomal position in megabases (Mbp) shown on the y-axis. For each chromosome pair, the color within the vertical bars indicates SNP density per 100 kb, ranging from low (purple) to high (yellow), black regions within the bars represent runs of homozygosity. Ribbons connecting the haplotypes illustrate structural annotations: syntenic regions (grey), inversions (orange), translocations (green), and duplications (blue).

### Differential methylation follows haplotype structure

We generated base-resolution 5mC methylation profiles for the diploid genome to investigate how the underlying haplotype structure influences epigenetic regulation. Comparing these profiles across major genomic features revealed a highly stable methylation landscape, with nearly identical methylation distributions between the two haplotypes in all three sequence contexts (CG, CHG, and CHH) (Figure 3A). The underlying patterns were canonical for a plant genome: TEs were hypermethylated, particularly in the CG ∼90% and CHG ∼70% contexts, whereas gene bodies were comparatively hypomethylated. To compare the epigenetic state of the genomic partitions, we constructed methylation metaprofiles across gene and TE bodies. This revealed a consistent hierarchy of methylation across all features and contexts: hemizygous regions consistently displayed the highest methylation levels, followed by homozygous regions, with heterozygous regions showing the lowest levels. For genes, all partitions exhibited the canonical M-shaped profile with sharp dips at the TSS and TES. The metaprofiles showed that TEs within hemizygous regions have markedly higher methylation levels in all three contexts, particularly CHG and CHH (Figure 3B). To understand the mechanism behind this, we investigated the relationship between TE age and methylation on a genome-wide scale. This analysis revealed that young, recently inserted TEs (<2% divergent) are the primary targets of dynamic silencing pathways, showing significantly higher CHG (Wilcoxon test, p < 2.2e-16) and CHH (Wilcoxon test, p < 2.2e-16) methylation than older TEs (>20% divergent) (Figure 3C). In contrast, CG methylation, associated with long-term repression, was uniformly high across TEs of all ages.

**Figure 3.**
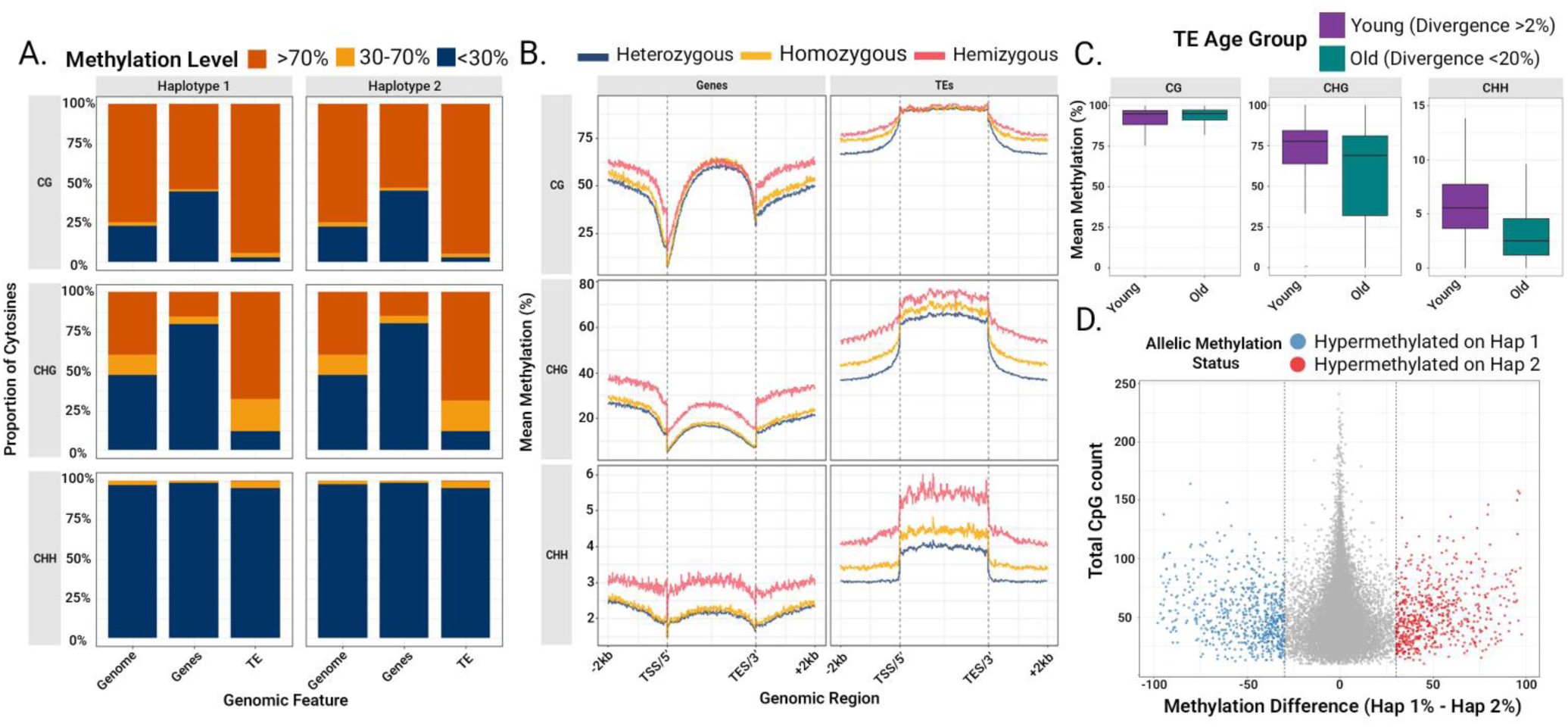
The Diploid Epigenomic Characterization of Pinot noir. **(a)** Global distribution of methylation levels across genomic features for Haplotype 1 and Haplotype 2. Proportions of cytosines are categorized as low (<30%), intermediate (30–70%), or highly (>70%) methylated within CG, CHG, and CHH contexts. **(b)** Metaplots displaying mean methylation levels across gene and TE bodies, bounded by transcription start/end sites (TSS/TES) and including 2 kb upstream and downstream flanking regions. Profiles are stratified by genomic partition: heterozygous (blue), homozygous (yellow), and hemizygous (red) regions. **(c)** Genome-wide comparison of mean methylation levels for young (<2% divergent) and old (>20% divergent) TEs **(d)** Scatter plot identifying Allele-Specific Methylation (ASM). Each point represents a one-to-one ortholog pair from heterozygous regions. The x-axis displays the difference in mean promoter CG methylation between alleles, while the y-axis represents the total CpG coverage (used as a proxy for confidence). Points falling beyond the dashed vertical lines (>30% difference) are classified as significant Asymmetrically Methylated Pairs (AMPs).

### Allelic methylation asymmetry is associated with local structural variation

Leveraging the diploid assembly and methylome, we next screened for allele-specific methylation (ASM). A genome-wide comparison of 16,754 one-to-one ortholog pairs in heterozygous regions identified 1,240 Asymmetrically Methylated Pairs (AMPs) where promoter mean methylation (CG context) differed by more than 30% between alleles (Figure 3D). These AMPs are distributed across all chromosomes (Figure S7) and show no bias toward either haplotype (588 hypermethylated on Haplotype 1 vs. 652 on Haplotype 2). Crucially, we found that this epigenetic asymmetry is strongly linked to local structural variation. A permutation test revealed that AMP promoters are significantly enriched for large InDels (>100bp) compared to the background of all gene promoters in heterozygous regions (Observed overlaps = 814; Empirical *p* < 0.001) (Figure S8), suggesting that local sequence divergence is a key factor associated with allele-specific epigenetic regulation. To establish a baseline epigenomic state for the reference genome, we classified all annotated genes based on their gene-body methylation (gbM) status, adapting the binomial test method of Takuno et al. (2017). Using a genic background CG methylation rate (*pCG*) of 0.535, we found that 31.4% of informative genes (15,537 of 49,454) were classified as body-methylated (BM), while 64.4% were undermethylated (UM) and 4.2% were intermediately methylated (IM) (Figure S9). When considering all 68,665 annotated genes, 22.6% were classified as gbM. The remaining genes were primarily unmethylated (46.4%) or had insufficient CG content (*n_CG_* < 20) for classification (28.0%)

### Rare Variants and Negative Selection shape the Somatic SNP Landscape

To define the nature and extent of somatic variation in grapevine, we performed Nanopore resequencing on 23 distinct Pinot noir clones. However, standard methods that map reads to a haploid reference are problematic for somatic variant calling in heterozygous species. By collapsing two distinct haplotypes into a single sequence, these methods often mistake allelic differences for somatic mutations or fail to detect real variants (Wang et al. 2024). While mapping to our diploid assembly minimized this bias, it created ambiguity for reads within extensive regions of homozygosity (RoH), leading to their exclusion by standard variant callers. To overcome this, we developed a novel ‘haplotype-masked’ reference that enforces unique read alignment in these tracts, enabling variant detection across the entire genome (see Supplementary Information). Our hybrid mapping strategy was highly effective. On average, 96.9% of reads from each clone mapped successfully to the masked diploid reference, achieving a mean depth across all samples of 26 X (Table S7). After applying a stringent filtering pipeline to variants called, we identified a final set of 67,277 high-quality somatic SNPs. The distribution of these variants was heavily skewed towards rarity, approximately 31% were singletons unique to a single clone (Figure 4A). Functionally, the landscape of somatic mutation is shaped by strong selective constraint. SNP density was strongly depleted in coding sequences compared to non-coding regions (Figure 4B). Intergenic regions exhibited the highest median density (7.1 SNPs per 100kb), followed by introns (2.6 SNPs per 100kb) and exons (1.2 SNPs per 100kb). Of the 948 variants within exons, the vast majority were either missense (575, 60.7%) or synonymous (356, 37.5%). High-impact mutations (e.g., nonsense) were exceedingly rare (32, 3.4%), consistent with purifying selection acting against deleterious somatic mutations in grapevine (Table S8).

**Figure 4.**
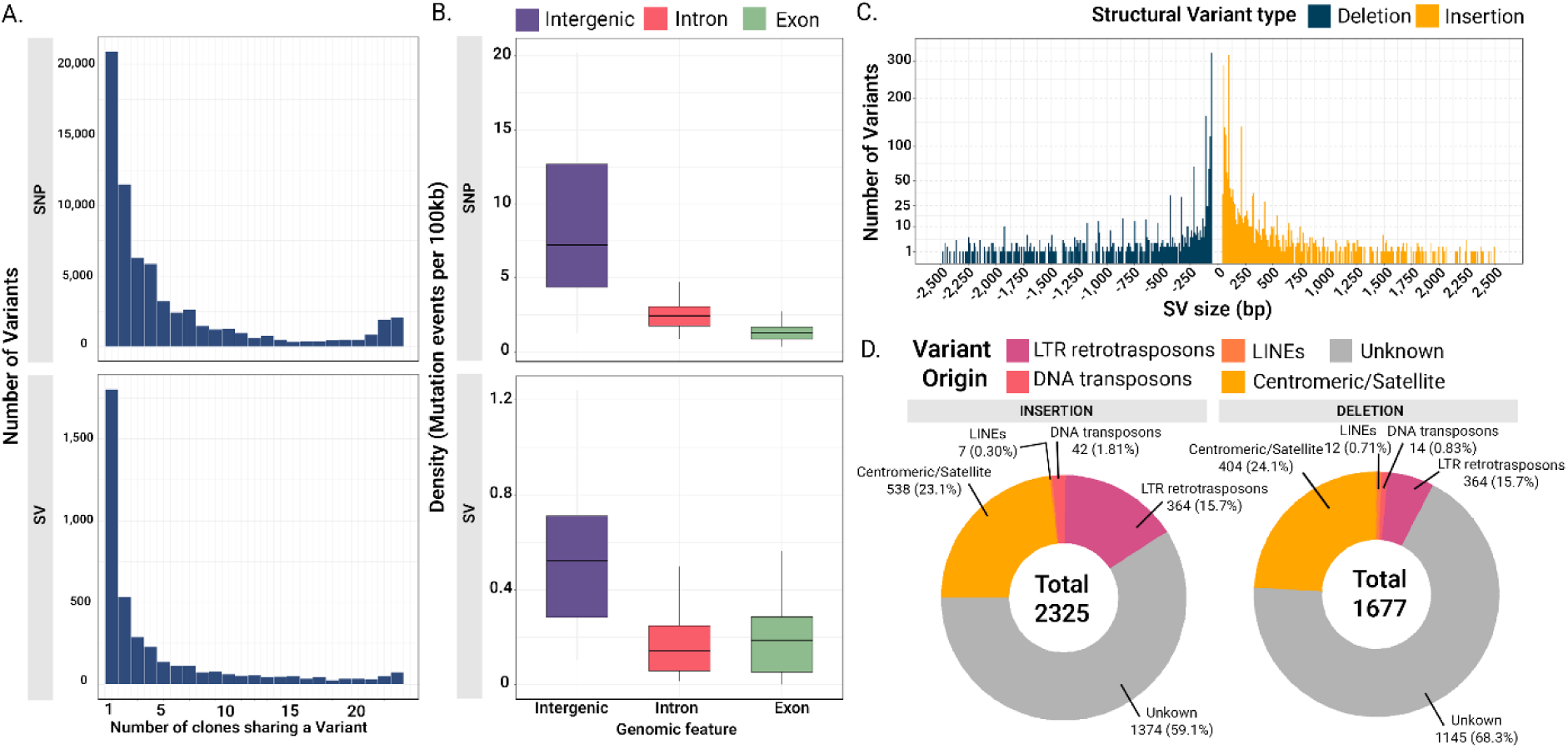
Comparative Analysis of Somatic SNP and SV. **(a)** Distribution of shared variants for SNPs (top) and SVs (bottom) **(b)** Density of somatic variants across genomic features for SNPs (top) and SVs (bottom). Boxplots show the distribution of variant event densities (events per 100kb) calculated for each of the 38 pseudo-chromosomes **(c)** Size spectrum of high-confidence insertions (yellow) and deletions (blue). Note that the y-axis is transformed to visualize lower-frequency large variants. **(d)** Mechanistic origin of insertions (left) and deletions (right). Donut charts show the proportion of SVs classified as originating from LTR retrotransposons, DNA transposons, LINEs, centromeric/satellite repeats, or of unknown origin.

### Repetitive DNA Shapes Somatic Structural Variant Formation Under Purifying Selection

Leveraging the ability of long reads to resolve complex events often missed by short-read approaches, we next characterized the landscape of structural variants larger than 50bp (SVs) using the long-read-aware caller Sniffles2 (Smolka et al. 2024). Our stringent filtering pipeline, which removed low-confidence calls and complex breakends (BNDs) prone to technical artifacts, yielded a final set of 4,037 high-confidence somatic SVs (>50 bp). This variant set was composed primarily of insertions (2,325; 57.6%) and deletions (1,677; 41.5%), with inversions (28; 0.7%) and duplications (7; 0.2%) being comparatively rare. Mirroring the pattern observed for SNPs, the SV frequency spectrum was heavily skewed towards rarity, with nearly half of all SVs (1,805; 44.7%) being private to a single clone (Figure 4A). Also consistent with SNPs, we observed a signal of purifying selection acting on SVs. The median SV density was highest in intergenic regions (0.52 SVs per 100kb) and was highly reduced in both introns (0.14 SVs per 100kb) and exons (0.18 SVs per 100kb), indicating selection against variants with deleterious potential (Figure 4B).

To investigate the mechanisms generating this variation, we analysed the SV size spectrum. The frequency of both insertions and deletions declined sharply with increasing variant size, consistent with purifying selection against large structural changes (Figure 4C). Interrupting this general decline were prominent, recurrent peaks corresponding to multiples of the 107 bp satellite repeat, the main component of grapevine centromeres (Shi et al. 2023). By classifying each SV by its size and sequence content, we formally quantified the contribution of different genomic features (Figure 4D). This analysis confirmed that centromeric satellite repeats are a major source of structural instability, accounting for 23.1% of insertions and 24.1% of deletions. LTR retrotransposons were the second-largest source, contributing substantially to insertions (15.7%) but less to deletions (6.1%). Together, these two classes of repetitive DNA accounted for a substantial fraction of all somatic insertions (38.8%) and deletions (30.2%) across the 23 Pinot noir clones.

### Extensive CG Gene-Body Methylation Variation Differentiates Clonal Genotypes

Leveraging the 23 clones base-resolution methylomes we performed a comparative analysis that identified 392,427 Differentially Methylated Cytosines (DMCs) across the panel. The majority of them are in the CG context (250,382), followed by CHG context (120,763) and a smaller fraction in the CHH context (21,282). The frequency spectra revealed a fundamental difference in stability: while non-CG variants were overwhelmingly rare, consistent with a transient, unstable state, CG-DMCs displayed a much broader distribution with many intermediate-frequency variants, indicating high mitotic stability (Figure 5A). In contrast to the patterns observed for genetic variants, the genomic distribution of DMCs was highly context-dependent (Figure 5B). DMCs in the CG context were significantly enriched within gene bodies, with the median density in exons (62.3 DMCs per 100kb) being substantially higher than in intergenic regions (20.4 per 100kb). The CHG context showed a different pattern, with DMCs being most abundant in introns (17.1 per 100kb) and exons (14.1 per 100kb), but depleted in intergenic regions (9.69 per 100kb). Finally, CHH DMCs were distributed more uniformly, with comparable median densities across intergenic (2.1 per 100kb), intronic (2.2 per 100kb), and exonic (2.3 per 100kb) features. We next explored whether these widespread CG DMCs translate to clone-specific changes in overall gene body methylation (gbM) status. Using the same gene-level binomial test as before, we assigned each gene in each clone to a BM, UM, or IM state. While the overall class composition was highly stable across clones (Figure 5C), we identified 3,716 genes (∼5.4% of all genes) that exhibited a clear gbM shift (switching between BM and UM states in different clones). A representative case is the gene *Vitis06g00265*, where a distinct clonal group is unmethylated (UM) across the coding sequence while several other clones are body-methylated (BM) (Figure 5D).

**Figure 5.**
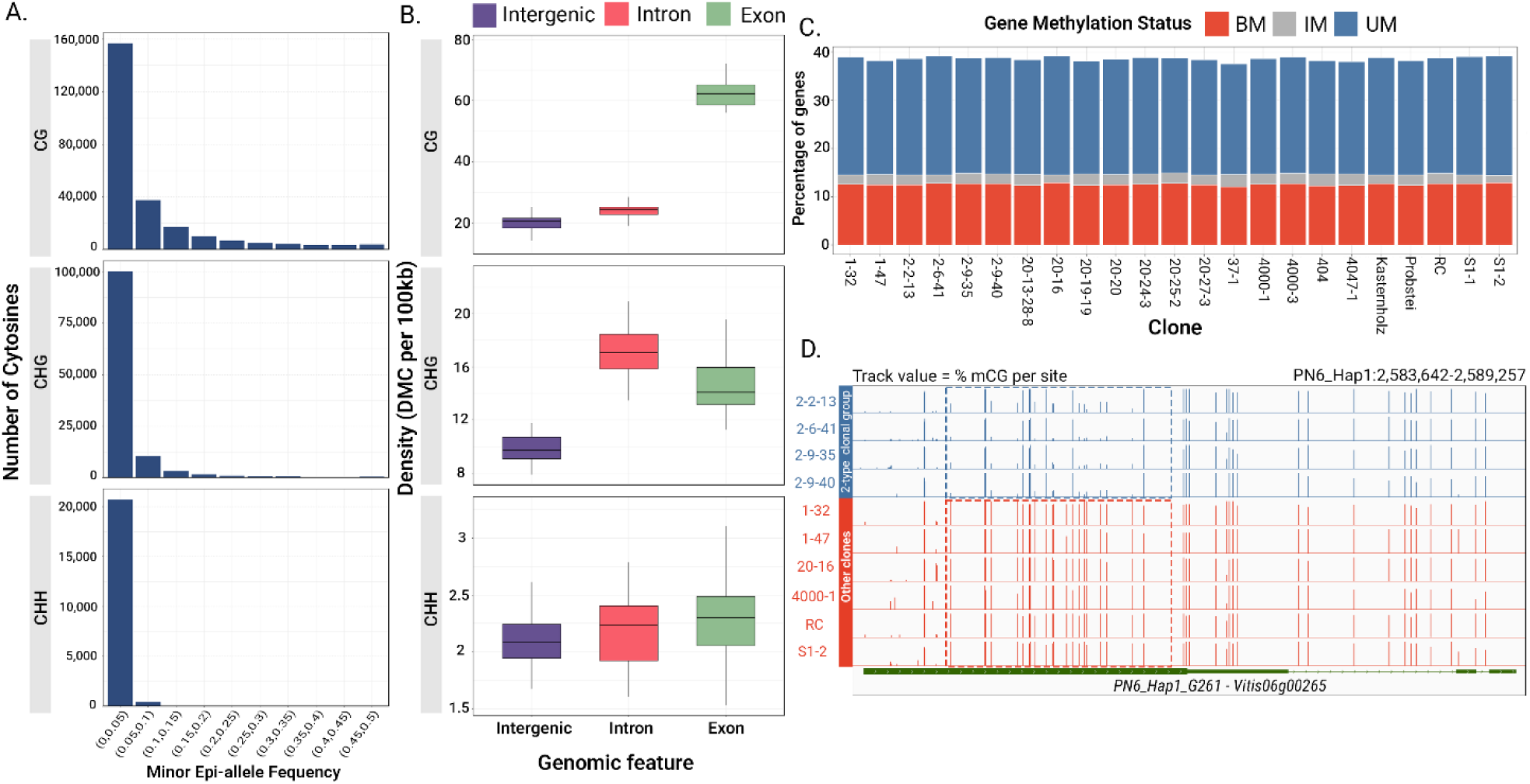
CG Methylation Varies Predominantly in Gene Bodies. **(a)** Minor Epi-allele Frequency (MEF) spectrum for DMCs in the CG, CHG, and CHH contexts. The x-axis represents the frequency of the minor methylation state (’0’ or ‘1’) among the 23 clones **(b)** Density of DMCs across genomic features, faceted by methylation context. Boxplots show the distribution of DMC event densities (events per 100kb) calculated for each of the 38 pseudo-chromosomes **(c)** Proportional distribution of gene body methylation (gbM) states across the 23 clones. For each clone, the bar shows the percentage of all informative genes classified as body-methylated (BM), intermediately methylated (IM), or unmethylated (UM) **(d)** Genome browser view of a representative gene (*Vitis06g00265*) exhibiting a clone-specific gbM shift. Each track represents the per-site CG methylation level for a single clone. Clones belonging to the “2-*” clonal group (blue tracks) are unmethylated across the gene body, while other clones (red tracks) are body-methylated.

### CG Epigenetic Signatures Mirror Genetic Lineages in Clonal Grapevines

Finally, we investigated whether the history of clonal propagation could be independently reconstructed from the patterns of genetic and epigenetic variation. For each data layer (SNPs, SVs, and DMCs in all three contexts), we compiled a clone-by-feature binary matrix, computed pairwise distances between all clones, and performed unsupervised hierarchical clustering.

As expected, the SNP-based phylogeny accurately resolved the known history of the collection, separating the two major clonal groups (20-Gm type and 2-Gm type) into distinct, well-supported clades (Figure 6A). Remarkably, the clustering based solely on CG methylation reproduces this genetic phylogeny with high fidelity (Figure 6A). This visual concordance was statistically confirmed by comparing the SNPs and CG-epigenetic distance matrices, which revealed an exceptionally strong positive correlation (Mantel statistic *r* = 0.96, *P* = 0.001) (Figure 6B). The major branches of the CG dendrogram are analogous to those of the SNP tree, grouping the same clones together with high confidence. This suggests that patterns of CG methylation are not random but stably inherited through vegetative propagation, accumulating over time to create a persistent epigenetic signature of clonal ancestry.

**Figure 6.**
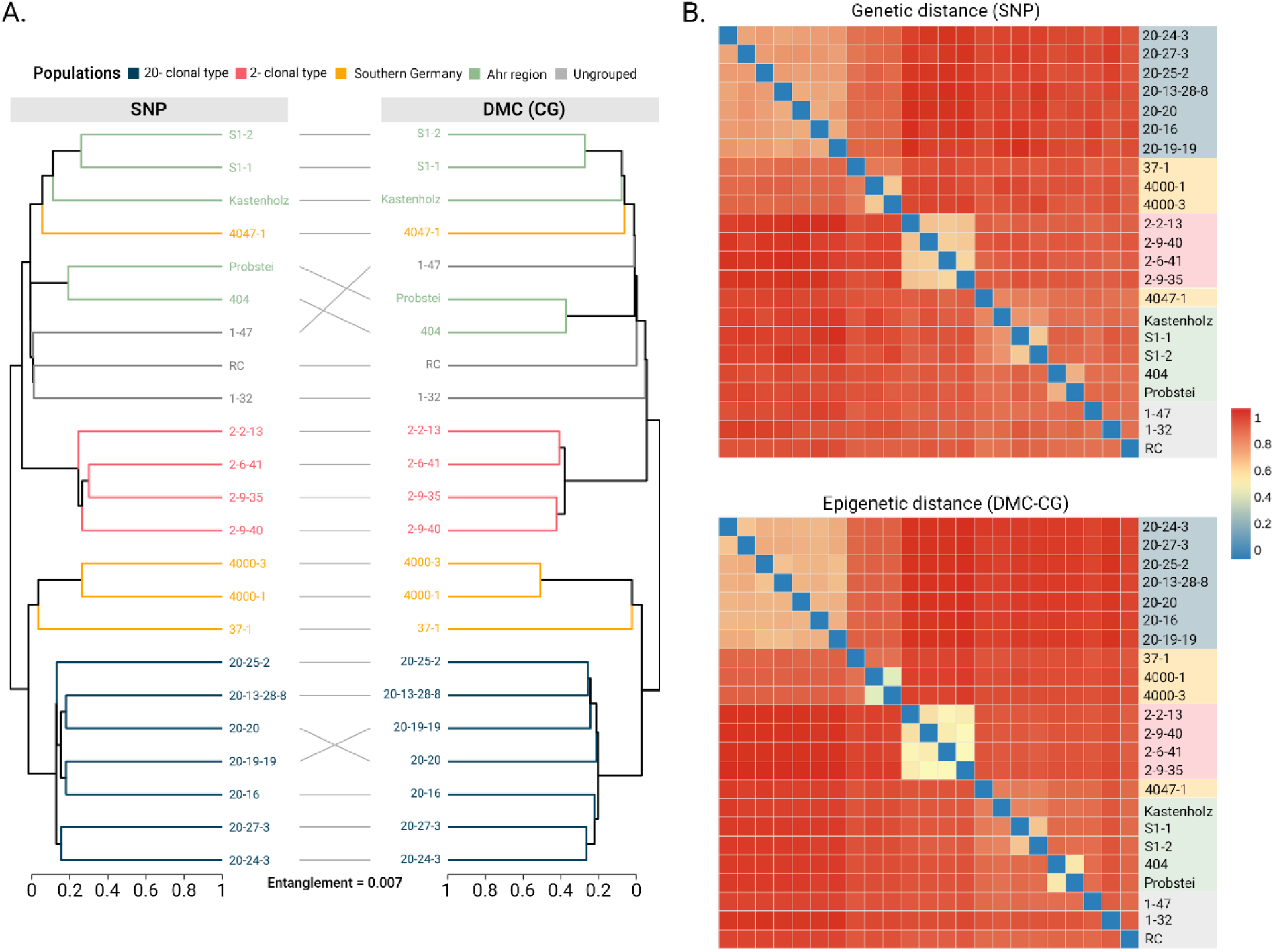
Genetic and Epigenetic Variation Reconstruct Clonal Propagation History. **(a)** Tanglegram comparing hierarchical clustering trees constructed from 67,277 genome-wide somatic SNPs (Left) and 250,382 somatic CG-context Differentially Methylated Cytosines (DMCs) (Right). Trees were inferred using distances derived from a Haploid Genetic Relationship Matrix (GRM) to account for the haploid somatic nature of the markers. Tips and connecting lines are colored by clonal groups defined a priori **(b)** Pairwise distance heatmaps for SNP (Top) and DMC-CG (Bottom) data *(D=1−GRM)*. To facilitate direct visual comparison, both heatmaps are sorted according to the Genetic (SNP) clustering order, and diagonal values were set to zero to normalize self-identity.

In contrast, the other data layers showed a weaker phylogenetic signal (Figure S11). Heatmaps based on SVs and CHG methylation only weakly recapitulated the major clonal groups, likely reflecting the high proportion of private, lineage-specific variants in these datasets. The CHH methylation landscape, however, appeared almost entirely random, with no discernible structure related to the known clonal relationships. This is consistent with the very low frequency and clone-specificity of CHH DMCs (Figure 5A). Collectively, these analyses suggest that clonal identity is defined by a combined mosaic of genetic mutations and stable CG epimutations, both of which can be used to accurately reconstruct the propagation history of grapevine lineages.

## Discussion

In clonally propagated perennials like grapevine, phenotypic diversity can arise from the accumulation of both somatic mutations and epigenetic changes (Berger et al. 2023). This is particularly relevant for grapevine cultivars which have been vegetatively propagated for centuries or longer. While both somatic mutations and epigenetic changes are known to contribute to clonal variability (Vondras et al. 2019, Varela et al. 2021, Xiao et al. 2025, Callipo et al. 2025), their relative importance, evolutionary dynamics and interplay remain poorly understood. This is largely because previous studies lacked the resolution to generate an integrated, genome-wide map of both variant types. Here, by generating a phased, diploid reference genome and leveraging nanopore long-read sequencing, we provide the first comprehensive view of the genetic and epigenetic mosaics that define clonal identity in Pinot noir.

While the recent T2T assembly of the inbred PN40024 (Shi et al. 2023) has been a major asset for grapevine geneticists, its homozygous nature fails to capture the complex heterozygosity of elite cultivars (Velt et al. 2023). Furthermore, although numerous high-quality diploid assemblies and pangenomes, including several for Pinot noir, have recently been published (Guo et al. 2025; Peng et al. 2025; Xiao et al. 2025; Liu et al. 2024), the substantial structural variation, often spanning hundreds of genes, found between individual clones (Alañón-Sánchez et al. 2025 *preprint*, Carbonell-Bejerano et al. 2017) warrants a specific reference for our Clonal population. Therefore, we generated a de novo assembly for the representative clone ‘20-13 Gm’. By sequencing native DNA, we simultaneously captured the genome and methylome, providing an integrated resource that complements existing assemblies.

Our diploid assembly reveals a genome defined by structural extremes. The identification of extensive runs of homozygosity (RoH) confirms a history of ancient inbreeding, a recurrent feature shared with other elite cultivars like Chardonnay (Roach et al. 2018), suggesting that ancestral inbreeding combined with long-term clonal propagation might be a recurrent genomic feature in historically important varieties. Furthermore, our analysis revealed the vast extent of the hemizygous fraction in the Pinot genome. Consistent with other clonally propagated species (Peng et al. 2025), these regions are primarily composed of repetitive elements and are strikingly gene-poor. This underscores hemizygosity as a fundamental and dynamic component of clonally propagated grapevines. Notably, we provide evidence that Transposable Element (TE) activity contributes substantially to diversification of these hemizygous regions. The TE sequence divergence profile within these tracts was bimodal, exhibiting a sharp peak of young elements with very low divergence. This pattern is a classic signature of a recent burst of transpositional activity, a method previously used to date major TE expansions and connect them with speciation in other clonally propagated fruit crops like apple (Daccord et al. 2017). This burst of young TEs, which was substantially weaker in heterozygous or homozygous regions, suggests that the ongoing, lineage-specific insertion of TEs, a known source of variation in grapevine (Zhou et al. 2019), creates novel genomic regions on one haplotype that are not present on the other.

Our integrated genome-methylome revealed high global methylation, particularly in the CG and CHG contexts, exceeding levels previously reported by standard bisulfite sequencing (Rodriguez-Izquierdo et al. 2024; Li et al. 2024; Peng et al. 2025; Zou et al. 2020). This increase is attributable to the reduced reference bias and superior mappability of long Nanopore reads in repetitive, TE-dense regions. The epigenomic landscape was globally stable, showing near-identical methylation distributions between the two haplotypes, a phenomenon already observed in cassava (Zhong et al. 2023) and figs (Usai et al. 2025). A clear hierarchy emerged upon partitioning by genomic structure: hemizygous regions consistently displayed the highest methylation levels (especially CHG and CHH). This elevated methylation correlates directly with the high density of young, recently inserted Transposable Elements (TEs) within these tracts. This observation is consistent with a model where the RNA-directed DNA Methylation (RdDM) pathway is the major force shaping the local epigenomic state of these novel genomic regions (Zhang et al. 2018).

Our diploid framework enabled the identification of thousands of genes subject to allele-specific methylation (ASM), where the two parental alleles of a gene exhibit different CG methylation patterns in their promoters (Figure 3D). We found that the promoters of these ASM genes were strongly enriched for large InDel variants. This underlying structural variation likely contributes to ASM by altering local chromatin accessibility or regulatory element spacing, facilitating the differential recruitment of methylation machinery. This provides molecular support for the view that structural variation is a major functional component in plants, capable of reshaping gene expression and modifying local regulatory environments (Yuan et al. 2021, Zhang et al. 2024).

Our integrated analysis of somatic variation confirms established principles of molecular evolution while providing new insight into the mechanisms that drive clonal divergence in grapevine. As expected, the genetic landscape is shaped by strong purifying selection. Both SNPs and SVs exhibit frequency spectra heavily skewed towards rarity, with a high proportion of private, clone-specific variants (Figure 4A, Figure 5A). This classic distribution, coupled with a significant depletion of variants from coding regions (Figure 4B), indicates strong selective constraint, consistent with prior work in clonally propagated grapevine (Cvijović et al. 2018, Vondras et al. 2019; Xiao et al. 2025).

Importantly, we found that structural variants arise largely in the highly repetitive fraction of the genome. The SV size spectrum was not random but highly structured, displaying prominent peaks corresponding precisely to multiples of the grapevine centromeric satellite repeats (Figure 4C). Mechanistic SV classification confirmed that these centromeric repeats and LTR retrotransposons represent the two largest identifiable sources of insertions and deletions (Figure 4D). These findings define a compartmentalized evolutionary landscape: purifying selection enforces strong constraint on coding regions, while highly unstable repetitive elements contribute disproportionally to structural plasticity.

Our study reveals that the epigenome is a rich and highly dynamic source of inter-clonal variation. We identified a conspicuously large amount of DMCs, with those in the CG context being the most numerous. The frequency spectra of DMCs revealed clear differences in epigenetic stability (Figure 5A). Non-CG (CHG and CHH) DMCs were almost exclusively rare, consistent with a transient and short-lived state (Kenchanmane Raju et al. 2019). Conversely, the spectrum for CG DMCs was much broader, likely reflecting the high fidelity of mCG maintenance through replication. Unlike genetic variants, CG-DMCs are not depleted from genes. Instead, they are significantly enriched within exons (Figure 5B), mirroring previous population-scale observations (Schmitz et al. 2013). This preferential targeting of gene bodies leads to widespread shifts in their methylation state. We identified thousands of genes that exhibit clear gbM shifts, switching between fully methylated and unmethylated states across clones. This finding, that thousands of genes exist as distinct “epialleles” within a clonal population, extends the concept of gbM-based variation (Shahzad et al. 2025) to a clonally propagated crop, where it may represent a mechanism for generating new phenotypic diversity in the absence of meiosis.

Our final analysis aimed to determine if clonal history could be reconstructed from different molecular layers. As expected, a phylogenetic tree derived from somatic SNPs accurately resolved known clonal groups, consistent with previous grapevine studies (Vondras et al. 2019; Gambino et al. 2017; Calderón et al. 2021). Notably, the CG epigenome provided a parallel and equally informative historical record. The high congruence between the SNP- and CG-DMC-based dendrograms (Figure 6) suggests that CG methylation patterns are not random but are mitotically inherited and stable across propagation cycles. This allows them to function as “biomarkers of origin” that reflect clonal ancestry, a phenomenon observed in other long-lived, clonally propagated organisms (Vanden Broeck et al. 2024; Yao et al. 2023). Together, these observations support the view that both somatic mutations and mitotically stable CG epialleles carry concordant, clock-like signals of clonal ancestry in perennial crops such as grapevine.

In summary, this work supports the view that clonal identity in grapevine reflects the combined contributions of both the genome and epigenome. We propose a model in which the coding genome is under strong selective constraint to preserve cultivar integrity, while the mitotically stable CG methylome represents a dynamic source of clonal variation. The discovery that stable CG epialleles track clonal history with the same phylogenetic resolution as SNPs provides a powerful new tool for understanding somatic evolution. This fundamentally broadens the paradigm of clonal variation, suggesting that vineyard phenotypic diversity is driven not only by rare genetic mutations but also by heritable changes in gene regulation. This perspective highlights the potential of “epi-breeding” for clonal improvement. By targeting stable epialleles through selection or direct methylation editing, it may be possible to fine-tune phenotypes without altering the underlying genotype. Such an approach offers a promising pathway to adapt elite cultivars like Pinot noir to emerging agricultural challenges while preserving their core genetic identity. Looking forward, future efforts should prioritize integrating high-throughput genetic and epigenetic fingerprinting with deep phenotyping across large clonal panels to precisely dissect the complex architecture of traits.

## Material and Methods

### Plant material

The Pinot clonal germplasm is maintained in a common experimental vineyard at the Department of Grapevine Breeding, Hochschule Geisenheim University (Geisenheim, Germany). This collection has been curated over several decades to preserve distinct agronomic and oenological phenotypes under homogeneous environmental conditions. The clone ‘20-13 Gm’ was selected for *de novo* reference genome assembly due to its widespread cultivation and economic importance in the region. Additionally, a diverse core panel of 23 clones was selected for re-sequencing to represent the breadth of phenotypic and geographical variation within the collection (Supplementary Information). For sampling, dormant woody cuttings were harvested from the field collection prior to winter pruning and rooted in perlite under greenhouse conditions. Upon establishment, young, expanding leaves (approximately 2–3 cm) were harvested from five independent ramets per clone. These tissues were pooled to generate a representative composite sample for each clone and immediately flash-frozen in liquid nitrogen for downstream DNA extraction.

### DNA extraction and sequencing

For the de novo assembly of the reference clone ‘20-13 Gm’, High-Molecular-Weight (HMW) DNA was isolated using the Nanobind PanDNA kit (PacBio). Two distinct sequencing libraries were generated to support a hybrid assembly strategy: an Oxford Nanopore ultra-long library, prepared using the Ultra-Long DNA Sequencing Kit (SQK-ULK114) and sequenced on two R10.4.1 flow cells (FLO-MIN114) using a GridION device; and a PacBio HiFi library, prepared and sequenced on a Revio system. For the re-sequencing panel of 23 clones, HMW DNA was extracted from pooled leaf tissue using the Genomic-tip 500/G kit (Qiagen) following the Nanopore’s suggested protocol for plant leaf tissue (Available at https://nanoporetech.com/document/extraction-method/fever-tree-gdna). To optimize read length distributions, DNA was mechanically sheared using a 26G needle and subsequently size-selected using the Short Read Eliminator XL kit (PacBio). DNA integrity and size distribution were verified by Pulsed-Field Gel Electrophoresis (PFGE), purity was assessed via NanoDrop spectrophotometry, and concentration was quantified using the Qubit dsDNA BR assay (Thermo Fisher). Sequencing libraries were prepared from 1.5µg of size selected DNA per genotype using the Ligation Sequencing Kit (SQK-LSK114) and sequenced on R10.4.1 flow cells (FLO-MIN114) using a GridION device.

### Basecalling and Methylation Calling

Raw Nanopore data were basecalled using Dorado v0.8.1 (https://github.com/nanoporetech/dorado) with the super-high-accuracy model (dna_r10.4.1_e8.2_400bps_sup@v5.0.0). Base modification probabilities for 5-methylcytosine (5mC) were generated concurrently during basecalling. These probabilities were subsequently aggregated into per-read and per-site methylation frequencies using modkit v0.4.3 (https://github.com/nanoporetech/modkit) (see Supplementary Methods for detailed parameters).

### Genome assembly and quality assessment

To generate a fully phased diploid assembly, PacBio HiFi reads and ONT ultra-long reads were co-assembled using hifiasm v0.21 (Cheng et al. 2021). The resulting primary contigs were scaffolded into chromosome-scale pseudomolecules using RagTag v2.1.0 (Alonge et al. 2022) utilizing as a guide the telomere-to-telomere PN40024 assembly (Shi et al. 2023). This process yielded two independent sets of 19 chromosomes representing the two haplotypes. Gene-space completeness was evaluated using BUSCO (Simão et al. 2015) against the eudicots_odb10 lineage dataset; Base-level accuracy and k-mer completeness were quantified using Merqury v1.3 (Rhie et al. 2020) comparing the assembly against a k-mer database derived from the high-fidelity PacBio reads. Structural integrity was assessed using CRAQ v1.0.9 (Li et al. 2023) for contiguity and structural accuracy metrics based on long-read support and reference consistency.

### Structural annotation

Protein-coding genes were annotated independently for each haplotype using a comprehensive pipeline that integrates ab initio prediction with homology-based evidence. First, *ab initio* gene models were generated using the deep-learning-based tool Helixer v0.2.0 (Holst et al. 2023). Simultaneously, high-quality reference gene models from PN40024.v4 (Velt et al., 2023) were mapped to the new assembly using Liftoff v1.6.3 (Shumate and Salzberg 2021). These two sets of evidence were then integrated, merged, and filtered using Mikado 2.3.4 (Venturini et al. 2018) to produce the final, high-confidence gene set.

Transposable elements (TEs) and other repetitive DNA were identified and annotated using the Extensive De novo TE Annotator (EDTA) pipeline v2.1.0 (Ou et al. 2019). The resulting TE annotation GFF3 file was used for all downstream analyses. The age of individual TE copies was estimated by calculating their sequence divergence from their consensus sequence. Divergence was calculated as *Divergence*(%) = 100 ∗ (1 − *identity*).

### Haplotype Comparison and Genomic Partitioning

To characterize the structural divergence between the two pseudo-haplotypes, a whole-genome alignment was performed using Minimap2 v2.28 (Li 2021). Structural rearrangements and syntenic regions were subsequently identified using SyRI v1.5 (Goel et al. 2019). The resulting structural annotations were visualized (As shown in Figure 2) using a customized implementation of PlotSR (Goel and Schneeberger 2022) available at https://github.com/HGU-Plant-Breeding/23_Pinot_Clones. Based on the SyRI output, the genome was partitioned in bins of 10kb into three primary structural states: Homozygous, defined as long, contiguous syntenic blocks (SYN) exhibiting near-zero sequence divergence (<5 SNPs/InDels per 10 kb).

Hemizygous, defined as a composite category encompassing non-aligned regions (NOTAL), large structural insertions/deletions (>1kb), and syntenic but highly diverged regions (HDR) where allelic correspondence is disrupted. Heterozygous, defined as syntenic blocks (SYN) containing a high density of small variants (see Supplementary Information).

The genomic composition of each partition was quantified using BEDTools v2.30 (Quinlan and Hall 2010). To assess allelic relationships, protein sequences from the primary transcripts of both haplotypes were analyzed using OrthoFinder (Emms and Kelly 2019). The resulting orthogroups were cross-referenced with the genomic partitions to determine gene copy number and synteny conservation.

Per-site methylation frequencies were extracted from the modkit output and converted into standard BedGraph format. To analyze epigenetic patterns across genomic features, methylation metaprofiles over gene and TE bodies were generated using deepTools v3.5.6 (Ramírez et al. 2014). To investigate the relationship between TE age and silencing, the mean methylation level for each individual TE copy was calculated using the bigWigAverageOverBed utility (Kent et al. 2010).

### Allelic methylation asymmetry (ASM) Analysis

To investigate Allele-Specific Methylation (ASM), one-to-one orthologs between the two haplotypes were first identified using OrthoFinder (Emms and Kelly 2019) and subsequently filtered to retain only those located within structurally defined heterozygous regions. Promoter regions were defined as the 2 kb sequence upstream of the Transcription Start Site (TSS). Mean promoter CG methylation levels were calculated using bigWigAverageOverBed (Kent et al. 2010). To ensure statistical robustness, analysis was restricted to promoters containing a minimum of 5 covered CpG sites on both alleles. Asymmetrically Methylated Pairs (AMPs) were defined as orthologs exhibiting an absolute difference in promoter methylation (Δ*m*CG) of ≥ 30%. AMP promoters were intersected with large InDels (≥100 bp) using BEDTools v2.30 (Quinlan and Hall 2010). In each iteration, a random set of promoters matching the sample size of the AMP set was drawn without replacement from the background of all analyzable heterozygous promoters. The empirical *p*-value was calculated as (*k* + 1)⁄(*N* + 1), where *k* is the number of random iterations exhibiting an overlap count equal to or greater than the observed value.

### Gene Body Methylation (gbM) Classification

Gene Body Methylation (gbM) status was classified for each gene in the genome assembly annotation. We applied the binomial test framework described by Takuno et al. (2017), calculating methylation over the Coding DNA Sequence (CDS) from start to stop codons. The null hypothesis (*p_CG_*) was defined as the average CG methylation rate of coding sequences for that specific clone. Only genes with ≥ 20 covered CG sites (*n_CG_*) were considered informative. Based on the binomial probability outcome, genes were classified into Body-Methylated (BM), Undermethylated (UM), or Intermediately Methylated (IM) states.

### Diploid-aware Read Mapping

To minimize reference bias and ensure accurate variant calling in a highly heterozygous genome, we employed a specialized “haplotype-masked” diploid mapping strategy.

First, a composite diploid reference was created by concatenating the two pseudo-haplotypes (PN_1 and PN_2). To resolve mapping ambiguity within Runs of Homozygosity (RoH), where identical sequences on both haplotypes typically cause reads to map with zero mapping quality (MAPQ=0), we masked the second haplotype. Using the coordinates of the homozygous regions previously defined, sequences corresponding to these tracts were systematically replaced with *’N’s* on the PN_2 haplotype using BEDTools maskfasta v2.30 (Quinlan and Hall 2010). This approach forces all reads originating from homozygous regions to align uniquely to PN_1, thereby rescuing their mapping quality while preserving diploid alignment in heterozygous regions (see Supplementary Information).

Nanopore reads from each of the 23 clones were aligned to this masked diploid reference using Minimap2 v2.28 (Li 2021) with the *map-ont* preset. The resulting alignments were sorted and indexed using Samtools v1.21 (Danecek et al. 2021).

### Variant Calling and Filtering

Somatic Single Nucleotide Polymorphisms (SNPs) were identified using a joint calling approach. Alignments from all 23 clones were processed simultaneously using bcftools mpileup with the Nanopore-specific configuration followed by bcftools call v1.15 (Danecek et al., 2021). The resulting multi-sample VCF was subjected to a stringent filtering pipeline to isolate high-confidence somatic variants. We retained only biallelic SNPs meeting the following criteria: QUAL > 10, INFO/MQ > 20, total depth (INFO/DP) between 115 and 690, and passing all internal bias tests (MQBZ > -2.5, BQBZ > -2.5, RPBZ between -2.5 and 2.5, and SCBZ < 5). Finally, to distinguish somatic mutations from fixed germline variants or systematic mapping artifacts, any locus where all 23 samples were genotyped as heterozygous was excluded.

Structural Variants (SVs) were identified using the population-level workflow of Sniffles2 (Smolka et al. 2024). First, SV candidates were quantified for each clone individually, generating. snf intermediates files. These were subsequently merged to produce a unified, multi-sample VCF. This raw callset was filtered using Bcftools to retain only high-confidence, resolvable SVs. We required variants to be marked as PASS in the FILTER field, QUAL > 30, be marked as PRECISE, have breakpoint and length standard deviations (STDEV_POS, STDEV_LEN) >50. Size selection was restricted to variants between 50 bp and 50 kb. Complex breakends (BNDs) were excluded to focus on discrete insertion, deletion, and inversion events.

### Genomic Distribution and Mechanistic Classification of Variants

To assess the functional distribution of somatic variants, the density of filtered SNPs and SVs (events per 100 kb) was calculated across exonic, intronic, and intergenic features for all 38 pseudo-chromosomes. Feature coordinates were derived from the consensus gene annotation, and densities were computed using custom scripts utilizing BEDTools v2.30 (Quinlan and Hall 2010). The allele frequency spectrum (AFS) was generated by quantifying the number of clones carrying the non-reference allele for each variant site.

To predict the functional impact of somatic point mutations, SNPs were annotated using SnpEff v5.2 (Cingolani et al., 2012). A custom database was constructed using the de novo assembly and the high-confidence gene annotation generated in this study. Variants were classified into impact categories (High, Moderate, Low, Modifier) based on their predicted effect on the coding sequence.

To elucidate the mechanisms driving structural variation, we analyzed the size spectrum and sequence content of SVs using a hierarchical classification strategy. First, the size distribution of insertions and deletions was visualized by plotting the SVLEN field from the filtered VCF. Second, variants were classified by origin: Deletions were classified as TE-related if they exhibited a > 80% reciprocal overlap with an annotated element from the EDTA library. Insertion sequences were extracted and queried against the EDTA TE library using BLASTN (Camacho et al., 2009). An insertion was classified as TE-derived if the best hit covered > 80% of the query length with > 80% identity. Variants not classified as TEs were checked for centromeric origin. SVs were classified as satellite-associated if their length corresponded to precise multiples (±1 bp) of known grapevine centromeres satellite repeat unit lengths (79, 107, 135, and 187 bp).

### Detection of Differentially Methylated Cytosines (DMCs)

To identify specific sites of epigenetic polymorphism, we developed a custom Python pipeline to process the per-site methylation tables generated by modkit. For each CpG site in each clone, the continuous methylation fraction was converted into a discrete binary state: “unmethylated” (< 30%) or “methylated” (> 70%). Sites exhibiting intermediate methylation (30-70%) or insufficient coverage (< 4x) were treated as missing data. The resulting binary matrix was filtered to retain only high-confidence DMCs with a successful call rate of ≥ 90% across the 23 clones. For each identified DMC, the Minor Epiallele Frequency (MEF) was calculated as the frequency of the least common binary state within the population. Genomic density (DMCs per 100kb) was calculated across exonic, intronic, and intergenic regions for all pseudo-chromosomes using the same annotation coordinates described above. Using the same binomial test as explained above, genes were classified per clone as BM/UM/IM within the CDS window. We required (*n_CG_* ≥ 20) sites per gene per clone; genes failing this were labelled as Low Coverage and excluded from class composition. The clone-specific pooled genic *p_CG_* served as the null. gbM-shift genes were those BM in ≥1 clone and UM in ≥1 other, the intermediate (IM) state was ignored for the purpose of defining binary shifts.

### Clonal lineage reconstruction from SNPs, SVs, and DMCs

To robustly reconstruct clonal phylogenies from somatic variation, we employed a quantitative genetics approach adapted for haplotype specific/somatic data. For each data layer (SNPs, SVs, and DMCs), genotypes were encoded as binary vectors (0=Reference/Unmethylated, 1=Alternative/Methylated). We calculated a Haploid Genomic Relationship Matrix (GRM) using a modified VanRaden method (VanRaden, 2008). Briefly, the genotype matrix *(M)* was centered by subtracting the allele frequency *(P)* to generate a centered matrix (*Z* = *M* − *P*). Missing data were handled via mean imputation on the centered matrix. The relationship matrix *(G)* was calculated as:

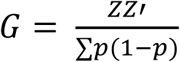

The scaling factor ∑*p*(1 − *p*) was used instead of the standard diploid 2∑*p*(1 − *p*) to correctly reflect the variance of somatic markers which exist as binary presence/absence states on a single haplotype lineage.

Clonal distances were derived from the relationship matrix as *D* = 1 − *G*. Hierarchical clustering was performed on these distance matrices using the UPGMA (Unweighted Pair Group Method with Arithmetic Mean) algorithm. To visually compare phylogenies, dendrograms were untangled and plotted as tanglegrams using the step2side algorithm implemented in the R package dendextend (Galili, 2015).

To statistically quantify the concordance between the genetic and epigenetic landscapes, a Mantel test was performed between the distance matrices using Spearman’s rank correlation with 9,999 permutations to determine significance, implemented using the R package vegan.

### Software and Customs Scripts

Primary bioinformatic processing was performed using the standard command-line tools detailed in the relevant methods sections. Custom scripts for downstream data processing, quantitative analysis, and statistical testing were developed in Python (v3.11.2) and R (v4.2.2). All final data visualizations were generated in R using the ggplot2 package (Wickham, 2016) and ggtree (Xu et al., 2022).

## Data and Code Availability

All raw sequencing data (PacBio HiFi and Oxford Nanopore) generated for the reference clone and the 23 clones sequencing panel have been deposited in the European Nucleotide Archive (ENA) under Project Accession PRJEB106155. The phased diploid genome assembly and corresponding functional annotations (GFF3) and methylation data (bedMethyl) are available at https://zenodo.org/records/18154549. All custom code and analysis pipelines developed for this study are publicly available on GitHub at https://github.com/HGU-Plant-Breeding/23_Pinot_Clones.

## Acknowledgments

Authors wish to thank Sara Riedel-Christ and Sonja Grundler for their assistance with greenhouse management and laboratory work at Hochschule Geisenheim University.

## Declaration of Generative AI and AI-assisted technologies in the writing process

During the preparation of this work the author(s) used Gemini pro 2.5 in order to improve the readability and the flow of the manuscript and to clean the code. After using this tool/service, the author(s) reviewed and edited the content as needed and take(s) full responsibility for the content of the publication.

## Authors’ contributions

K.P.V-F., M.S. and P.C. conceived and designed the study. P.C. and M.S. performed the laboratory work for DNA extraction and sequencing. P.C. performed the bioinformatic analyses. P.C. developed the custom analysis scripts and created all figures. M.S. and H.R. interpreted results, provided critical suggestions on the methods and participated in the discussion. K.P.V.F. supervised the project and acquired funding. P.C. wrote the original manuscript draft. All authors contributed to the discussion of the results and the review and editing of the final manuscript.

## Publication bibliography

Noelia Alañón-Sánchez, Yolanda Ferradás, Ilja Bezrukov et al. A complex reciprocal translocation underlies reduced bunch compactness in a grapevine somatic variant, 06 October 2025, PREPRINT (Version 1) available at Research Square [10.21203/rs.3.rs-7594260/v1]

Alonge, Michael; Lebeigle, Ludivine; Kirsche, Melanie; Jenike, Katie; Ou, Shujun; Aganezov, Sergey et al. (2022): Automated assembly scaffolding using RagTag elevates a new tomato system for high-throughput genome editing. In Genome Biology 23 (1), p. 258. DOI: 10.1186/s13059-022-02823-7.

Alston, Julian M.; Sambucci, Olena (2019): Grapes in the World Economy. In Dario Cantu, M. Andrew Walker (Eds.): The Grape Genome. Cham: Springer International Publishing, pp. 1–24.

Berger, Margot M. J.; Stammitti, Linda; Carrillo, Natalia; Blancquaert, Erna; Rubio, Bernadette; Teyssier, Emeline; Gallusci, Philippe (2023): Epigenetics: an innovative lever for grapevine breeding in times of climatic changes: This article is published in cooperation with the 22nd GiESCO International Meeting, hosted by Cornell University in Ithaca, NY, July 17-21, 2023. In OENO One 57 (2), pp. 265–282. DOI: 10.20870/oeno-one.2023.57.2.7405.

Calderón, Luciano; Mauri, Nuria; Muñoz, Claudio; Carbonell-Bejerano, Pablo; Bree, Laura; Bergamin, Daniel et al. (2021): Whole genome resequencing and custom genotyping unveil clonal lineages in ‘Malbec’ grapevines (Vitis vinifera L.). In Scientific Reports 11 (1), p. 7775. DOI: 10.1038/s41598-021-87445-y.

Callipo, Paolo; Schmidt, Maximilian; Strack, Timo; Robinson, Hannah; Vasudevan, Akshaya; Voss-Fels, Kai P. (2025): Harnessing clonal diversity in grapevine: from genomic insights to modern breeding applications. In Theoretical and Applied Genetics 138 (8), p. 196. DOI: 10.1007/s00122-025-04986-w.

Carbonell-Bejerano, Pablo; Royo, Carolina; Torres-Pérez, Rafael; Grimplet, Jérôme; Fernandez, Lucie; Franco-Zorrilla, José Manuel et al. (2017): Catastrophic Unbalanced Genome Rearrangements Cause Somatic Loss of Berry Color in Grapevine. In Plant physiology 175 (2), pp. 786–801. DOI: 10.1104/pp.17.00715.

Carvalho, Luisa C.; Pinto, Teresa; Costa, Joaquim Miguel; Martins, Antero; Amâncio, Sara; Gonçalves, Elsa (2025): Intra-Varietal Variability for Abiotic Stress Tolerance Traits in the Grapevine Variety Arinto. In Plants 14 (16). DOI: 10.3390/plants14162480.

Cheng, Haoyu; Concepcion, Gregory T.; Feng, Xiaowen; Zhang, Haowen; Li, Heng (2021): Haplotype-resolved de novo assembly using phased assembly graphs with hifiasm. In Nature Methods 18 (2), pp. 170–175. DOI: 10.1038/s41592-020-01056-5.

Cingolani, Pablo; Platts, Adrian; Wang, Le L.; Coon, M.; Nguyen, T.; Wang, L.; Land, S. J.; Lu, X.; Ruden, Douglas M. (2012): A program for annotating and predicting the effects of single nucleotide polymorphisms, SnpEff: SNPs in the genome of Drosophila melanogaster strain w1118; iso-2; iso-3. In Fly 6 (2), pp. 80–92. DOI: 10.4161/fly.19695.

Cvijović, Ivana; Good, Benjamin H.; Desai, Michael M. (2018): The Effect of Strong Purifying Selection on Genetic Diversity. In Genetics 209 (4), pp. 1235–1278. DOI: 10.1534/genetics.118.301058.

Daccord, Nicolas; Celton, Jean-Marc; Linsmith, Gareth; Becker, Claude; Choisne, Nathalie; Schijlen, Elio et al. (2017): High-quality de novo assembly of the apple genome and methylome dynamics of early fruit development. In Nature Genetics 49 (7), pp. 1099–1106. DOI: 10.1038/ng.3886.

Danecek, Petr; Bonfield, James K.; Liddle, Jennifer; Marshall, John; Ohan, Valeriu; Pollard, Martin O. et al. (2021): Twelve years of SAMtools and BCFtools. In Gigascience 10 (2), giab008. DOI: 10.1093/gigascience/giab008.

Dong, Yang; Duan, Shengchang; Xia, Qiuju; Liang, Zhenchang; Dong, Xiao; Margaryan, Kristine et al. (2023): Dual domestications and origin of traits in grapevine evolution. In Science 379 (6635), pp. 892–901. DOI: 10.1126/science.add8655.

Emms, David M.; Kelly, Steven (2019): OrthoFinder: phylogenetic orthology inference for comparative genomics. In Genome Biology 20 (1), p. 238. DOI: 10.1186/s13059-019-1832-y.

Gambino, Giorgio; Dal Molin, Alessandra; Boccacci, Paolo; Minio, Andrea; Chitarra, Walter; Avanzato, Carla Giuseppina et al. (2017): Whole-genome sequencing and SNV genotyping of ‘Nebbiolo’ (Vitis vinifera L.) clones. In Scientific Reports 7 (1), p. 17294. DOI: 10.1038/s41598-017-17405-y.

Goel, Manish; Schneeberger, Korbinian (2022): plotsr: visualizing structural similarities and rearrangements between multiple genomes. In Bioinformatics 38 (10), pp. 2922–2926. DOI: 10.1093/bioinformatics/btac196.

Goel, Manish; Sun, Hequan; Jiao, Wen-Biao; Schneeberger, Korbinian (2019): SyRI: finding genomic rearrangements and local sequence differences from whole-genome assemblies. In Genome Biology 20 (1), p. 277. DOI: 10.1186/s13059-019-1911-0.

Guo, Li; Wang, Xiangfeng; Ayhan, Dilay Hazal; Rhaman, Mohammad Saidur; Yan, Ming; Jiang, Jianfu et al. (2025): Super pangenome of Vitis empowers identification of downy mildew resistance genes for grapevine improvement. In Nature Genetics 57 (3), pp. 741–753. DOI: 10.1038/s41588-025-02111-7.

Guo, Ying; Feng, Yang-Fan; Yang, Gang-Gui; Jia, Yan; He, Jie; Wu, Ze-Yu et al. (2024): Allele-specific DNA methylation and gene expression during shoot organogenesis in tissue culture of hybrid poplar. In Hortic Res 11 (3), uhae027. DOI: 10.1093/hr/uhae027.

Holst, Felix; Bolger, Anthony; Günther, Christopher; Maß, Janina; Triesch, Sebastian; Kindel, Felicitas et al. (2023): Helixer de novo Prediction of Primary Eukaryotic Gene Models Combining Deep Learning and a Hidden Markov Model. In bioRxiv. DOI: 10.1101/2023.02.06.527280.

Jaillon, Olivier; Aury, Jean-Marc; Noel, Benjamin; Policriti, Alberto; Clepet, Christian; Casagrande, Alberto et al. (2007): The grapevine genome sequence suggests ancestral hexaploidization in major angiosperm phyla. In Nature 449 (7161), pp. 463–467. DOI: 10.1038/nature06148.

Kenchanmane Raju, Sunil K.; Ritter, Eleanore Jeanne; Niederhuth, Chad E. (2019): Establishment, maintenance, and biological roles of non-CG methylation in plants. In Essays in biochemistry 63 (6), pp. 743–755. DOI: 10.1042/EBC20190032.

Kent, W. James; Zweig, Ann S.; Barber, G.; Hinrichs, A. S.; Karolchik, D. (2010): BigWig and BigBed: enabling browsing of large distributed datasets. In Bioinformatics 26 (17), pp. 2204–2207. DOI: 10.1093/bioinformatics/btq351.

Lacombe, Thierry; Boursiquot, Jean-Michel; Laucou, Valérie; Di Vecchi-Staraz, Manuel; Péros, Jean-Pierre; This, Patrice (2013): Large-scale parentage analysis in an extended set of grapevine cultivars (Vitis vinifera L.). In Theoretical and Applied Genetics 126 (2), pp. 401–414. DOI: 10.1007/s00122-012-1988-2.

Li, Heng (2021) New strategies to improve minimap2 alignment accuracy, Bioinformatics, Volume 37, Issue 23, Pages 4572–4574 DOI: 10.1093/bioinformatics/btab705

Li, Ao; Wang, Fengxia; Ding, Tingting; Li, Ke; Liu, Huiping; Zhang, Qingtian et al. (2024): Genome-wide DNA methylation dynamics and RNA-seq analysis during grape (cv. ‘Cabernet Franc’) skin coloration. In Genomics 116 (2), p. 110810. DOI: 10.1016/j.ygeno.2024.110810.

Li, Kunpeng; Xu, Peng; Wang, Jinpeng; Yi, Xin; Jiao, Yuannian (2023): Identification of errors in draft genome assemblies at single-nucleotide resolution for quality assessment and improvement. In Nature Communications 14 (1), p. 6556. DOI: 10.1038/s41467-023-42336-w.

Liu, Zhongjie; Wang, Nan; Su, Ying; Long, Qiming; Peng, Yanling; Shangguan, Lingfei et al. (2024): Grapevine pangenome facilitates trait genetics and genomic breeding. In Nature Genetics 56 (12), pp. 2804–2814. DOI: 10.1038/s41588-024-01967-5.

McGovern, Patrick E. (2003): Ancient Wine. The Search for the Origins of Viniculture. Princeton: Princeton University Press.

Ocaña, Juan; Walter, Bernard; Schellenbaum, Paul (2013): Stable MSAP Markers for the Distinction of Vitis vinifera cv Pinot Noir Clones. In Molecular Biotechnology 55 (3), pp. 236–248. DOI: 10.1007/s12033-013-9675-3.

OIV (2025): The State of the World Vine and Wine Sector in 2024. Dijon. Available online at https://www.oiv.int/sites/default/files/2025-04/OIV-State_of_the_World_Vine-and-Wine-Sector-in-2024.pdf, checked on 9/19/2025.

Ou, Shujun; Su, Weija; Liao, Yi; Chougule, Kapeel; Agda, Jireh R. A.; Hellinga, Adam J. et al. (2019): Benchmarking transposable element annotation methods for creation of a streamlined, comprehensive pipeline. In Genome Biology 20 (1), p. 275. DOI: 10.1186/s13059-019-1905-y.

Pelsy, F. (2010): Molecular and cellular mechanisms of diversity within grapevine varieties. In Heredity 104 (4), pp. 331–340. DOI: 10.1038/hdy.2009.161.

Peng, Yanling; Wang, Yiwen; Liu, Yuting; Fang, Xinyue; Cheng, Lin; Long, Qiming et al. (2025): The genomic and epigenomic landscapes of hemizygous genes across crops with contrasting reproductive systems. In Proceedings of the National Academy of Sciences 122 (6), e2422487122. DOI: 10.1073/pnas.2422487122.

Portu, Javier; Baroja, Elisa; Rivacoba, Luis; Martínez, Juana; Ibáñez, Sergio; Tello, Javier (2024): Evaluation of the intra-varietal diversity of ‘Tempranillo Tinto’ clones prospected in the demarcated winemaking region of Rioja (Spain). In Scientia Horticulturae 329, p. 113015. DOI: 10.1016/j.scienta.2024.113015.

Quinlan, Aaron R.; Hall, Ira M. (2010): BEDTools: a flexible suite of utilities for comparing genomic features. In Bioinformatics 26 (6), pp. 841–842. DOI: 10.1093/bioinformatics/btq033.

Ramírez, Fidel; Dündar, Friederike; Diehl, Sarah; Grüning, Björn A.; Manke, Thomas (2014): deepTools: a flexible platform for exploring deep-sequencing data. In Nucleic Acids Res 42 (W1), W187–W191. DOI: 10.1093/nar/gku365.

Ramos-Madrigal, Jazmín; Runge, Anne Kathrine Wiborg; Bouby, Laurent; Lacombe, Thierry; Samaniego Castruita, José Alfredo; Adam-Blondon, Anne-Françoise et al. (2019): Palaeogenomic insights into the origins of French grapevine diversity. In Nature Plants 5 (6), pp. 595–603. DOI: 10.1038/s41477-019-0437-5.

Regner, Ferdinand; Stadlbauer, Alexandra; Eisenheld, Cornelia; Kaserer, Herwlg (2000): Genetic Relationships Among Pinots and Related Cultivars. In American Journal of Enology and Viticulture 51 (1), pp. 7–14. DOI: 10.5344/ajev.2000.51.1.7.

Rhie, Arang; Walenz, Brian P.; Koren, Sergey; Phillippy, Adam M. (2020): Merqury: reference-free quality, completeness, and phasing assessment for genome assemblies. In Genome Biology 21 (1), p. 245. DOI: 10.1186/s13059-020-02134-9.

Roach, Michael J.; Johnson, Daniel L.; Bohlmann, Joerg; van Vuuren, Hennie J. J.; Jones, Steven J. M.; Pretorius, Isak S. et al. (2018): Population sequencing reveals clonal diversity and ancestral inbreeding in the grapevine cultivar Chardonnay. In PLOS Genetics 14 (11), e1007807. DOI: 10.1371/journal.pgen.1007807.

Rodriguez-Izquierdo, Alberto; Carrasco, David; Anand, Lakshay; Magnani, Roberta; Catarecha, Pablo; Arroyo-Garcia, Rosa; Rodriguez Lopez, Carlos M. (2024): Epigenetic differences between wild and cultivated grapevines highlight the contribution of DNA methylation during crop domestication. In BMC Plant Biology 24 (1), p. 504. DOI: 10.1186/s12870-024-05197-z.

Schmidt, Maximilian; Strack, Timo; Andrews, Haylie; Hickey, Lee T.; Crisp, Peter A.; Voss-Fels, Kai P. (2025): A new climate for genomic and epigenomic innovation in grapevine. In Molecular Horticulture 5 (1), p. 44. DOI: 10.1186/s43897-025-00171-1.

Schmitz, Robert J.; Schultz, Matthew D.; Urich, Mark A.; Nery, Joseph R.; Pelizzola, Mattia; Libiger, Ondrej et al. (2013): Patterns of population epigenomic diversity. In Nature 495 (7440), pp. 193–198. DOI: 10.1038/nature11968.

Shahzad, Zaigham; Hollwey, Elizabeth; Moore, Jonathan D.; Choi, Jaemyung; Cassin-Ross, Gaëlle; Rouached, Hatem et al. (2025): Gene body methylation regulates gene expression and mediates phenotypic diversity in natural Arabidopsis populations. In Nature Plants 11, 2084–2099. DOI: 10.1038/s41477-025-02108-4.

Shi, Xiaoya; Cao, Shuo; Wang, Xu; Huang, Siyang; Wang, Yue; Liu, Zhongjie et al. (2023): The complete reference genome for grapevine (Vitis vinifera L.) genetics and breeding. In Hortic Res 10 (5), uhad061. DOI: 10.1093/hr/uhad061.

Shumate, Alaina; Salzberg, Steven L. (2021): Liftoff: accurate mapping of gene annotations. In Bioinformatics 37 (12), pp. 1639–1643. DOI: 10.1093/bioinformatics/btaa1016.

Simão, Felipe A.; Waterhouse, Robert M.; Ioannidis, Panagiotis; Kriventseva, Evgenia V.; Zdobnov, Evgeny M. (2015): BUSCO: assessing genome assembly and annotation completeness with single-copy orthologs. In Bioinformatics 31 (19), pp. 3210–3212. DOI: 10.1093/bioinformatics/btv351.

Smolka, Moritz; Paulin, Luis F.; Grochowski, Christopher M.; Horner, Dominic W.; Mahmoud, Medhat; Behera, Sairam et al. (2024): Detection of mosaic and population-level structural variants with Sniffles2. In Nature Biotechnology 42 (10), pp. 1571–1580. DOI: 10.1038/s41587-023-02024-y.

Takuno, Shohei; Seymour, Danelle K.; Gaut, Brandon S. (2017): The Evolutionary Dynamics of Orthologs That Shift in Gene Body Methylation between Arabidopsis Species. In Mol Biol Evol 34 (6), pp. 1479–1491. DOI: 10.1093/molbev/msx099.

This, Patrice; Lacombe, Thierry; Thomas, Mark R. (2006): Historical origins and genetic diversity of wine grapes. In Trends in genetics 22 (9), pp. 511–519. DOI: 10.1016/j.tig.2006.07.008.

Urra, Claudio; Sanhueza, Dayan; Pavez, Catalina; Tapia, Patricio; Núñez-Lillo, Gerardo; Minio, Andrea et al. (2023): Identification of grapevine clones via high-throughput amplicon sequencing: a proof-of-concept study. In G3: Genes,Genomes,Genetics 13 (9), jkad145. DOI: 10.1093/g3journal/jkad145.

Usai, Gabriele; Giordani, Tommaso; Vangelisti, Alberto; Castellacci, Marco; Simoni, Samuel; Bosi, Emanuele et al. (2025): Haplotype-resolved genome assembly of Ficus carica L. reveals allele-specific expression in the fruit. In Plant J 121 (4), e70012. DOI: 10.1111/tpj.70012.

van Houten, Silvina; Muñoz, Claudio; Bree, Laura; Bergamín, Daniel; Sola, Cristobal; Lijavetzky, Diego (2020): Natural Genetic Variation for Grapevine Phenology as a Tool for Climate Change Adaptation. In Applied Sciences 10 (16). DOI: 10.3390/app10165573.

an Vanden Broeck; Meese, Tim; Verschelde, Pieter; Cox, Karen; Heinze, Berthold; Deforce, Dieter et al. (2024): Genome-wide methylome stability and parental effects in the worldwide distributed Lombardy poplar. In BMC Biology 22 (1), p. 30. DOI: 10.1186/s12915-024-01816-1.

Varela, Anabella; Ibañez, Verónica N.; Alonso, Rodrigo; Zavallo, Diego; Asurmendi, Sebastián; Gomez Talquenca, Sebastián et al. (2021): Vineyard environments influence Malbec grapevine phenotypic traits and DNA methylation patterns in a clone-dependent way. In Plant Cell Reports 40 (1), pp. 111–125. DOI: 10.1007/s00299-020-02617-w.

Velasco, Riccardo; Zharkikh, Andrey; Troggio, Michela; Cartwright, Dustin A.; Cestaro, Alessandro; Pruss, Dmitry et al. (2007): A High Quality Draft Consensus Sequence of the Genome of a Heterozygous Grapevine Variety. In PLOS ONE 2 (12), e1326. DOI: 10.1371/journal.pone.0001326.

Velt, Amandine; Frommer, Bianca; Blanc, Sophie; Holtgräwe, Daniela; Duchêne, Éric; Dumas, Vincent et al. (2023): An improved reference of the grapevine genome reasserts the origin of the PN40024 highly homozygous genotype. In G3 Genes|Genomes|Genetics 13 (5), jkad067. DOI: 10.1093/g3journal/jkad067.

Venturini, Luca; Caim, Shabhonam; Kaithakottil, Gemy George; Mapleson, Daniel Lee; Swarbreck, David (2018): Leveraging multiple transcriptome assembly methods for improved gene structure annotation. In Gigascience 7 (8), giy093. DOI: 10.1093/gigascience/giy093.

Vezzulli, Silvia; Leonardelli, Lorena; Malossini, Umberto; Stefanini, Marco; Velasco, Riccardo; Moser, Claudio (2012): Pinot blanc and Pinot gris arose as independent somatic mutations of Pinot noir. In J Exp Bot 63 (18), pp. 6359–6369. DOI: 10.1093/jxb/ers290.

Vondras, Amanda M.; Minio, Andrea; Blanco-Ulate, Barbara; Figueroa-Balderas, Rosa; Penn, Michael A.; Zhou, Yongfeng et al. (2019): The genomic diversification of grapevine clones. In BMC Genomics 20 (1), p. 972. DOI: 10.1186/s12864-019-6211-2.

Wang, Nan; Chen, Peng; Xu, Yuanyuan; Guo, Lingxia; Li, Xianxin; Yi, Hualin et al. (2024): Phased genomics reveals hidden somatic mutations and provides insight into fruit development in sweet orange. In Hortic Res 11 (2), uhad268. DOI: 10.1093/hr/uhad268.

Wickham, Hadley (2016): ggplot2: Elegant Graphics for Data Analysis. Springer-Verlag New York

Xiao, Hua; Wang, Yue; Liu, Wenwen; Shi, Xiaoya; Huang, Siyang; Cao, Shuo et al. (2025): Impacts of reproductive systems on grapevine genome and breeding. In Nature Communications 16 (1), p. 2031. DOI: 10.1038/s41467-025-56817-7.

Xu, Shuangbin; Li, Lin; Luo, Xiao; Chen, Meijun; Tang, Wenli; Zhan, Li; Dai, Zehan; Lam, Tommy T.; Guan, Yi; Yu, Guangchuang (2022): Ggtree: A serialized data object for visualization of a phylogenetic tree and annotation data. In iMeta 1, e56. DOI: 10.1002/imt2.56.

Yao, N.; Zhang, Z.; Yu, L.; Hazarika, R.; Yu, C.; Jang, H. et al. (2023): An evolutionary epigenetic clock in plants. In Science 381 (6665), pp. 1440–1445. DOI: 10.1126/science.adh9443.

Yuan, Yuxuan; Bayer, Philipp E.; Batley, Jacqueline; Edwards, David (2021): Current status of structural variation studies in plants. In Plant Biotechnol J 19 (11), pp. 2153–2163. DOI: 10.1111/pbi.13646.

Zhang, Huiming; Lang, Zhaobo; Zhu, Jian-Kang (2018): Dynamics and function of DNA methylation in plants. In Nature Reviews Molecular Cell Biology 19 (8), pp. 489–506. DOI: 10.1038/s41580-018-0016-z.

Zhang, Yuanyuan; Yang, Zhiquan; He, Yizhou; Liu, Dongxu; Liu, Yueying; Liang, Congyuan et al. (2024): Structural variation reshapes population gene expression and trait variation in 2,105 Brassica napus accessions. In Nature Genetics 56 (11), pp. 2538–2550. DOI: 10.1038/s41588-024-01957-7.

Zhong, Zhenhui; Feng, Suhua; Mansfeld, Ben N.; Ke, Yunqing; Qi, Weihong; Lim, Yi-Wen et al. (2023): Haplotype-resolved DNA methylome of African cassava genome. In Plant Biotechnol J 21 (2), pp. 247–249. DOI: 10.1111/pbi.13955.

Zhou, Yongfeng; Massonnet, Mélanie; Sanjak, Jaleal S.; Cantu, Dario; Gaut, Brandon S. (2017): Evolutionary genomics of grape (Vitis vinifera ssp. vinifera) domestication. In Proceedings of the National Academy of Sciences 114 (44), pp. 11715–11720. DOI: 10.1073/pnas.1709257114.

Zhou, Yongfeng; Minio, Andrea; Massonnet, Mélanie; Solares, Edwin; Lv, Yuanda; Beridze, Tengiz et al. (2019): The population genetics of structural variants in grapevine domestication. In Nature Plants 5 (9), pp. 965–979. DOI: 10.1038/s41477-019-0507-8.

Zou, Luming; Liu, Wenwen; Zhang, Zhan; Edwards, Everard J.; Gathunga, Elias Kirabi; Fan, Peige et al. (2020): Gene body demethylation increases expression and is associated with self-pruning during grape genome duplication. In Hortic Res 7 (1), p. 84. DOI: 10.1038/s41438-020-0303-7.

