## Supplementary Information for "A dual genomic-epigenomic map of clonal evolution in grapevine"

#### Supplementary Methods

##### Plant Material and Core Panel Selection

The Pinot clonal germplasm is maintained at the Department of Plant Breeding, Hochschule Geisenheim University (Geisenheim, Germany). This extensive collection comprises over 200 unique accessions, sourced between the late 20th and early 21st centuries from major viticultural regions across Germany and Europe. Accessions were introduced into the common experimental vineyard at various time points; consequently, while all plants are maintained under identical environmental conditions, some accessions have undergone multiple vegetative propagation cycles at the Geisenheim site.

From this wider collection, a reference clone (20-13 Gm) and a core panel of 23 clones were selected to maximize genetic and phenotypic diversity. Selection was guided by two primary criteria:

Geographical Diversity: The panel captures a broad cross-section of the German viticultural landscape, representing all major growing regions. This local diversity is complemented by reference accessions from France and Switzerland to provide a broader European genetic context.

Phenotypic Contrast: Clones were chosen to cover the full spectrum of cluster architecture phenotypes characteristic of Pinot noir, ranging from loose-clustered morphotypes (e.g., the '1-Gm' type) to highly compact forms (e.g., clone '4047-1'). Additionally, variation in vegetative growth habits, specifically the inclusion of erect-growing types (e.g., the '2-Gm' type), and differences in ripening kinetics were key selection factors.

##### Genome Assembly and Scaffolding Parameters

PacBio HiFi reads and Oxford Nanopore ultra-long reads (filtered for length >50 kb) were co-assembled using hifiasm v0.21 in diploid mode. The specific execution command utilized the ultra-long integration module to resolve complex repetitive regions:

```
hifiasm -o ./PN-20-13.asm --telo-m TTTAGGG -s 0.3 --dual-scaf -t 128 --ul 20-13_ONT_50KB.fastq 20-13_HiFi.fastq
```

The resulting phased assembly graphs (.gfa) were converted to FASTA format, and the contiguity was visually assessed using dotplots generated by Minimap2 and D-GENIES.

The primary contigs for each haplotype were scaffolded against the Pinot Noir assembly published by (Liu et al. 2024) using RagTag v2.1.0 in scaffold mode, the command used for Haplotype 1 (representative of both haplotypes) was:

```
ragtag.py scaffold T2T.fasta PN-20_13_Hap_1.fa -q 20 -i 0.5 -a 0.5 -C -o PN-20_13_Hap_1_to_T2T -t 4
```

The resulting scaffolds were visually inspected using dotplots generated by Minimap2 and D-GENIES.

#### Custom PlotSR Visualization

To generate the integrated genomic visualization (Figure 2), we utilized the PlotSR tool (Goel and Schneeberger, 2022). We cloned the original source code from the official repository (https://github.com/schneebergerlab/plotsr) and extended the core Python library by implementing custom plotting functions.

Specifically, we implemented a custom function to parse window-based SNP density data and render it as a continuous color-gradient heatmap directly on the chromosome bars. This modification allows for the simultaneous visualization of structural rearrangements (ribbons) and local genetic diversity (heatmap) in a single track.

The modified source code is available at: https://github.com/HGU-Plant-Breeding/23_Pinot_Clones/tree/main/1_Genome_Assembly/plotsr_modified.

#### Genomic Partitioning Strategy and Rationale

To dissect the structural architecture of the diploid genome, we employed a hierarchical subtraction strategy to partition the genome into three mutually exclusive structural states. The partition was defined sequentially as follows:

*Definition of Hemizygous Regions*

Firstly, we defined regions lacking a linear allelic counterpart. This category included:

NOTAL: Regions classified as "Not Aligned" by SyRI (sequence unique to one haplotype).

Large SVs: Insertions and Deletions defined by SyRI with a length >1 kb

HDR: "Highly Diverged Regions" of at least 1kb where synteny exists but sequence identity is too low to support read alignment.

A size threshold of 1 kb was chosen to distinguish macro-structural variation from local "micro-hemizygosity" (e.g., small promoter indels). This cutoff ensures the capture of complete functional units, such as intact Transposable Elements (TEs) or whole gene models, representing true structural turnover rather than minor polymorphism.

*Definition of Homozygous Regions (Runs of Homozygosity)*

Next, we identified regions of ancient inbreeding. We selected contiguous blocks classified as Syntenic (SYN) by SyRI that exhibited a variant density below 5 small variants (SNPs + InDels) per 10kb. This specific threshold was empirically determined following extensive visual inspection of read alignments using the Integrative Genomics Viewer (IGV). We observed that within genuine homozygous tracts, the variant density is typically near-zero. Therefore, a cutoff of < 0.0005 variants/bp serves as a conservative upper bound, effectively accommodating rare sequencing errors or isolated somatic mutations while robustly excluding the heterozygous background.

*Definition of Heterozygous Regions*

The Heterozygous partition was defined as the genomic remainder:

$$Heterozygous = Total\ Genome - (Homozygous + Hemizygous)$$

By defining this category via subtraction, we capture the "core" genome regions that are syntenic and alignable (unlike Hemizygous) but possess standard inter-allelic variation (unlike Homozygous). This ensures that the three categories are mathematically exhaustive and non-overlapping.

This logic is implemented in the custom Python script *classify_genome.py*, which takes the SyRI output and chromosome sizes as input and outputs a color-coded BED file representing the three partition states.

#### Gene Body Methylation (gbM) Classification Algorithm

To classify the methylation status of gene bodies, we adopted the statistical framework established by Takuno et al. (2017). This method employs a binomial test to determine whether the methylation level of a specific coding sequence significantly deviates from the background methylation rate of the genome (or clone).

We developed two custom Python scripts to implement the Takuno et al. (2017) framework specifically for high-throughput Nanopore data. Both scripts apply identical statistical parameters (α=0.05, Min Sites=20, Min Coverage=60%) to ensure results are strictly comparable.

First, to define the baseline epigenomic landscape of the reference clone, we utilized a single-sample implementation (*classify_gbm_single.py*). This script calculates the global background methylation rate ($p_{CG}$) specifically for the '20-13 Gm' diploid assembly and tests every annotated gene against this static null hypothesis, generating the baseline statistics reported in the Results.

To extend this analysis to the clonal population, we developed a matrix-mode wrapper (*classify_gbm_matrix.py*) tailored for comparative epigenomics. Rather than applying a fixed reference background, this tool iterates the binomial test across all 23 re-sequenced clones, calculating a dynamic null hypothesis ($p_{CG}$) specific to each individual sample. This sample-specific normalization is critical for distinguishing true biological variation from technical noise arising from slight inter-sample differences in global methylation levels. The pipeline aggregates these individual tests to produce a unified Gene × Clone classification matrix, which serves as the foundation for identifying "Epiallele Shifts" genes that exhibit stable state switching (e.g., BM to UM) between different clonal lineages.

#### Diploid-Aware Mapping and Haplotype Masking Strategy

To minimize reference bias, our goal was to align reads to a fully diploid reference containing both pseudo-haplotypes (PN_1 and PN_2). However, initial benchmarks revealed a critical limitation: in regions of high sequence identity, specifically the extensive Runs of Homozygosity (RoH) identified by SyRI, sequencing reads aligned with equal probability to both haplotypes. This "mapping ambiguity" resulted in a Mapping Quality (MAPQ) score of zero, causing these reads to be systematically discarded by downstream variant callers and creating artificial blind spots in the analysis.

To resolve this, we implemented a "Haplotype-Masked" reference strategy that strictly defines the alignment behavior based on genomic structure:

*Heterozygous Regions (Natural Separation):* In structurally divergent or heterozygous regions, both haplotypes are retained in the reference. Reads naturally align to their haplotype of origin based on sequence specificity, preserving phasing information.

*Homozygous Regions (Forced Alignment):* Within the RoH coordinates defined by SyRI, we selectively masked the sequence of the secondary haplotype (PN_2) with 'N's. This masking effectively removes the ambiguity: reads originating from these homozygous tracts are physically unable to align to the masked PN_2 sequence and are therefore forced to align uniquely to the corresponding locus on PN_1.

This strategy successfully "rescues" the mapping quality of reads in homozygous regions, raising their MAPQ scores to high confidence levels. Consequently, this allows for the detection of rare somatic mutations even within the highly conserved inbred regions of the genome, while simultaneously maintaining full diploid resolution in the heterozygous core.

#### Methylation Data Processing and DMC Identification Strategy

To identify Differentially Methylated Cytosines (DMCs) with high confidence, we developed a custom bioinformatics framework designed to transition from raw probabilistic signals to robust, binary epigenetic genotypes. This pipeline was structured to address specific biological properties of plant methylation and technical characteristics of Nanopore sequencing.

*Strand Merging and Signal Integration*

We began by extracting base-modification probabilities using modkit pileup using the command:

```
modkit pileup ${SAMPLE}.bam ${SAMPLE}_all_c.bed --motif CG 0 --motif CHG 0 --motif CHH 0 --ignore h --ref ${REF} --threads 4
```

However, treating forward and reverse strands independently can lead to reduced effective coverage and stochastic noise. We implemented a strand-merging strategy to increase data robustness. Using custom Python scripts, we merged symmetrical cytosines, adjacent CpGs on opposite strands and symmetrical CHG motifs. For these sites, the methylation level was recalculated as the coverage-weighted average of both strands. This step effectively doubled the read depth for symmetric contexts, providing a more reliable estimate of the local methylation state. Conversely, CHH methylation, which is established de novo and is typically asymmetric, was processed in a strand-specific manner to preserve biological accuracy.

*Discretization and Ambiguity buffering*

A critical challenge in methylation analysis is distinguishing true epigenetic states from technical noise or cell-type heterogeneity. To address this, we eschewed continuous value comparisons in favor of a ternary discretization strategy. We defined strict high-confidence thresholds: sites with >70% methylation were classified as Methylated (1), and sites with <30% as Unmethylated (0). Crucially, we treated the intermediate interval (30–70%) as "Ambiguous" rather than forcing a binary call. This "buffer zone" ensures that loci exhibiting heterogeneous methylation patterns or noisy signal accumulation are excluded from binary variant calling, thereby minimizing false positives. Additionally, a minimum coverage threshold of 4× was enforced to ensure sufficient statistical power for these calls.

*Population Matrix Construction and Filtering*

To scale this analysis to the population level, we employed a parallelized workflow to aggregate individual sample data into a unified Site × Clone matrix. This matrix was then subjected to stringent quality filtering to define the final set of DMCs. To prevent missing data from confounding distance metrics, we required a Call Rate ≥90%, retaining only sites that were confidently genotyped (0 or 1) in at least 21 of the 23 clones. Finally, to identify somatic polymorphisms, we filtered for sites with a Minor Epiallele Frequency (MEF) >0, retaining only those loci where at least one clone exhibited a high-confidence methylation state change relative to the population consensus.

### Supplementary Figures

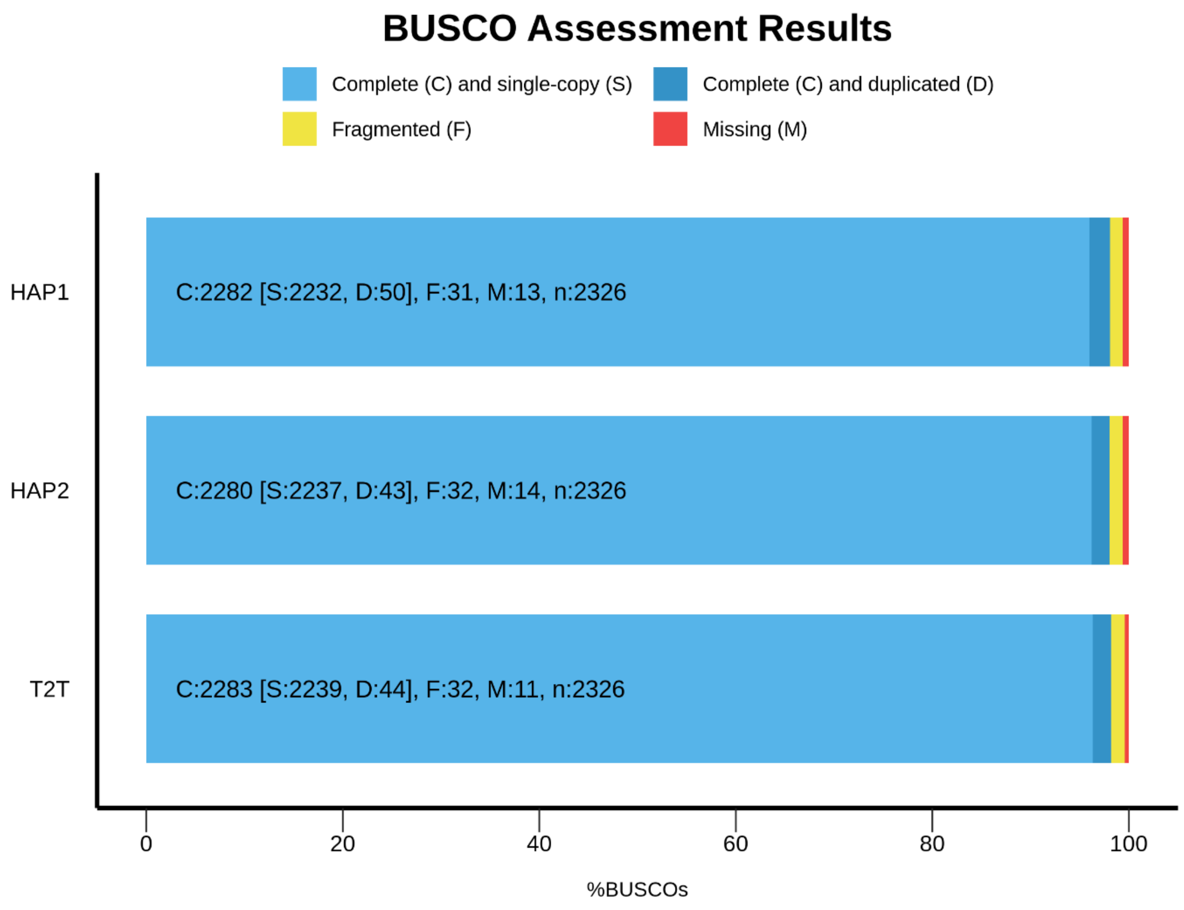


**Figure S1. BUSCO Assessment of Genome Assembly Completeness**

Bar chart showing the completeness of the two pseudo-haplotypes (HAP1, HAP2) and the PN40024 T2T reference assembly based on the BUSCO (Benchmarking Universal Single-Copy Orthologs). The bars show the percentage of BUSCOs that were found as complete and single-copy (S), complete and duplicated (D), fragmented (F), or missing (M).

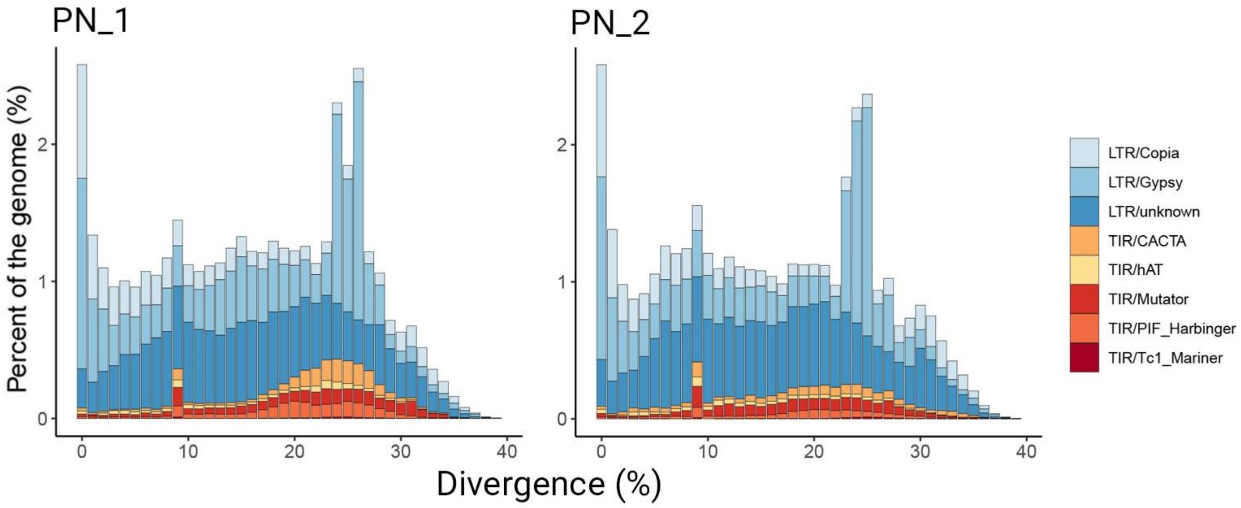


**Figure S2. Transposable Element Divergence Profiles for Each Haplotype.**

Histograms showing the genomic landscape of transposable elements (TEs) for the two pseudo-haplotypes, PN_1 and PN_2. The y-axis represents the percentage of the genome each category occupies. The x-axis represents the sequence divergence (%) of individual TE copies from their consensus sequence, a proxy for TE age. Colors indicate the major superfamilies of LTR and TIR retrotransposons.

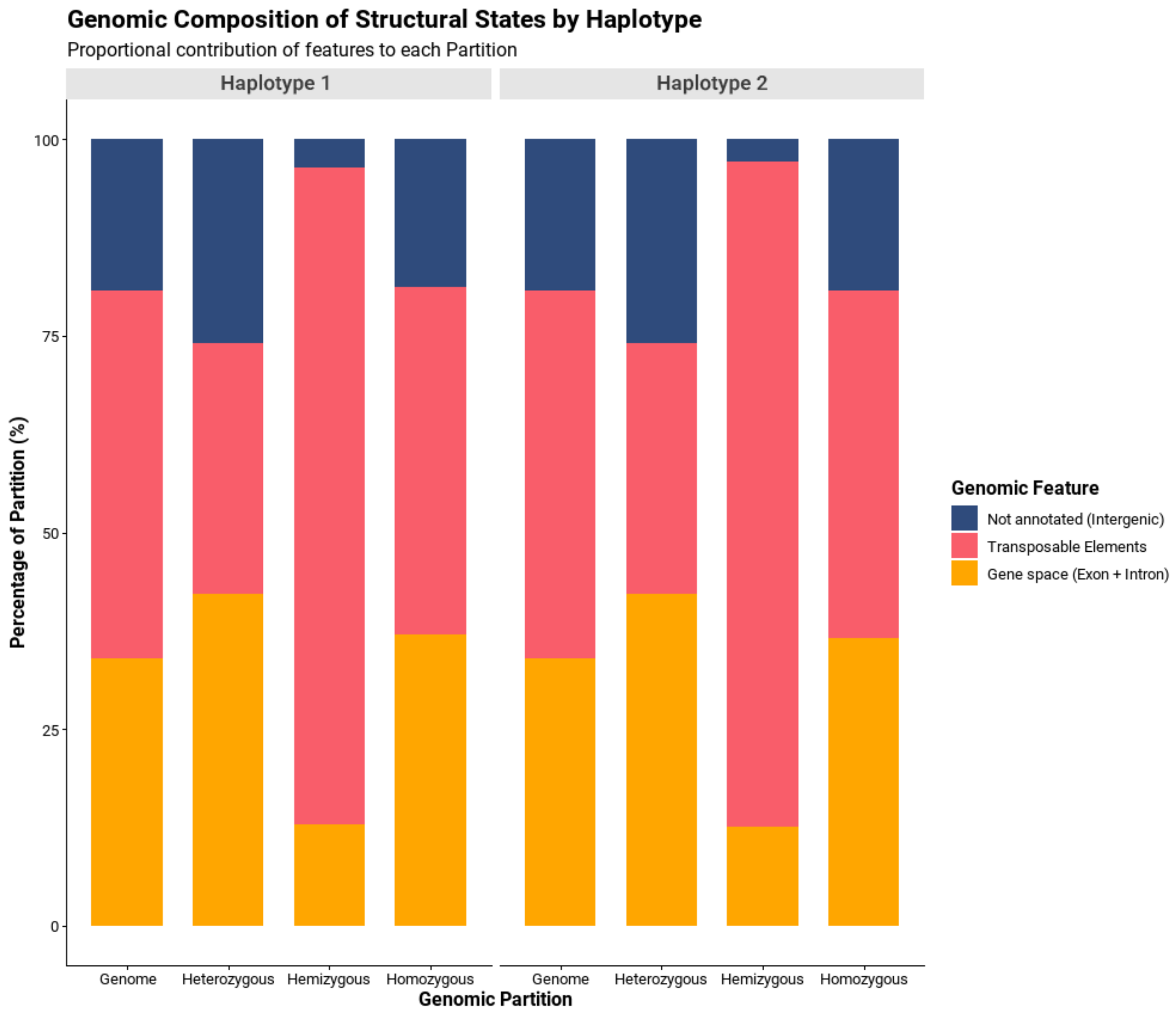


#### Figure S3. Genomic Composition of Structural States

Stacked bar charts showing the proportional contribution of different genomic features to the overall genome and to each of the three primary structural partitions (Heterozygous, Hemizygous, Homozygous) in both haplotypes.

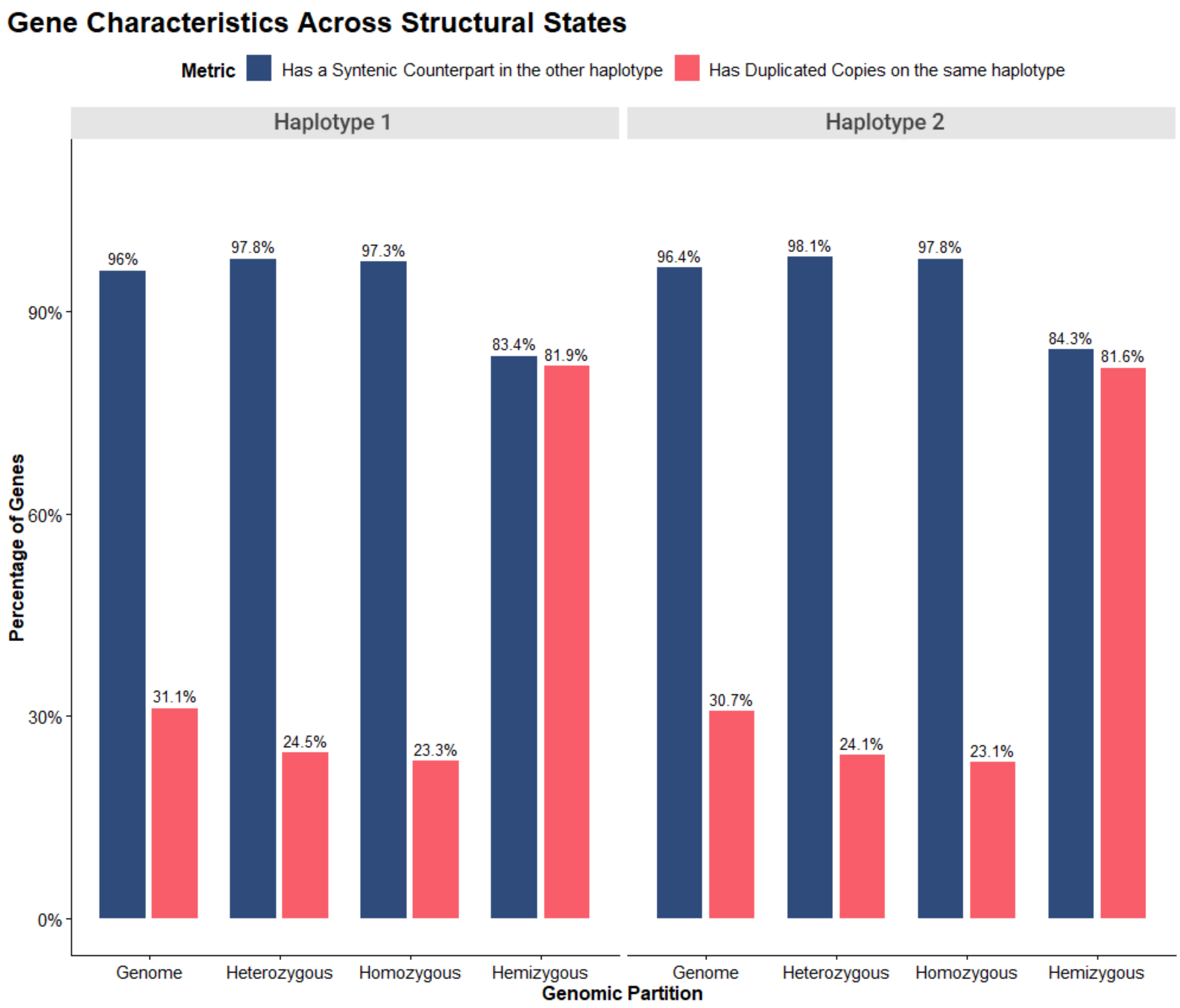


#### Figure S4. Gene Characteristics Across Structural States

Bar charts comparing key characteristics of genes located within the three genomic partitions. For each partition and each haplotype, the blue bar shows the percentage of genes that have a direct syntenic counterpart on the opposing haplotype. The red bar shows the percentage of genes that have at least one duplicated copy within the same haplotype.

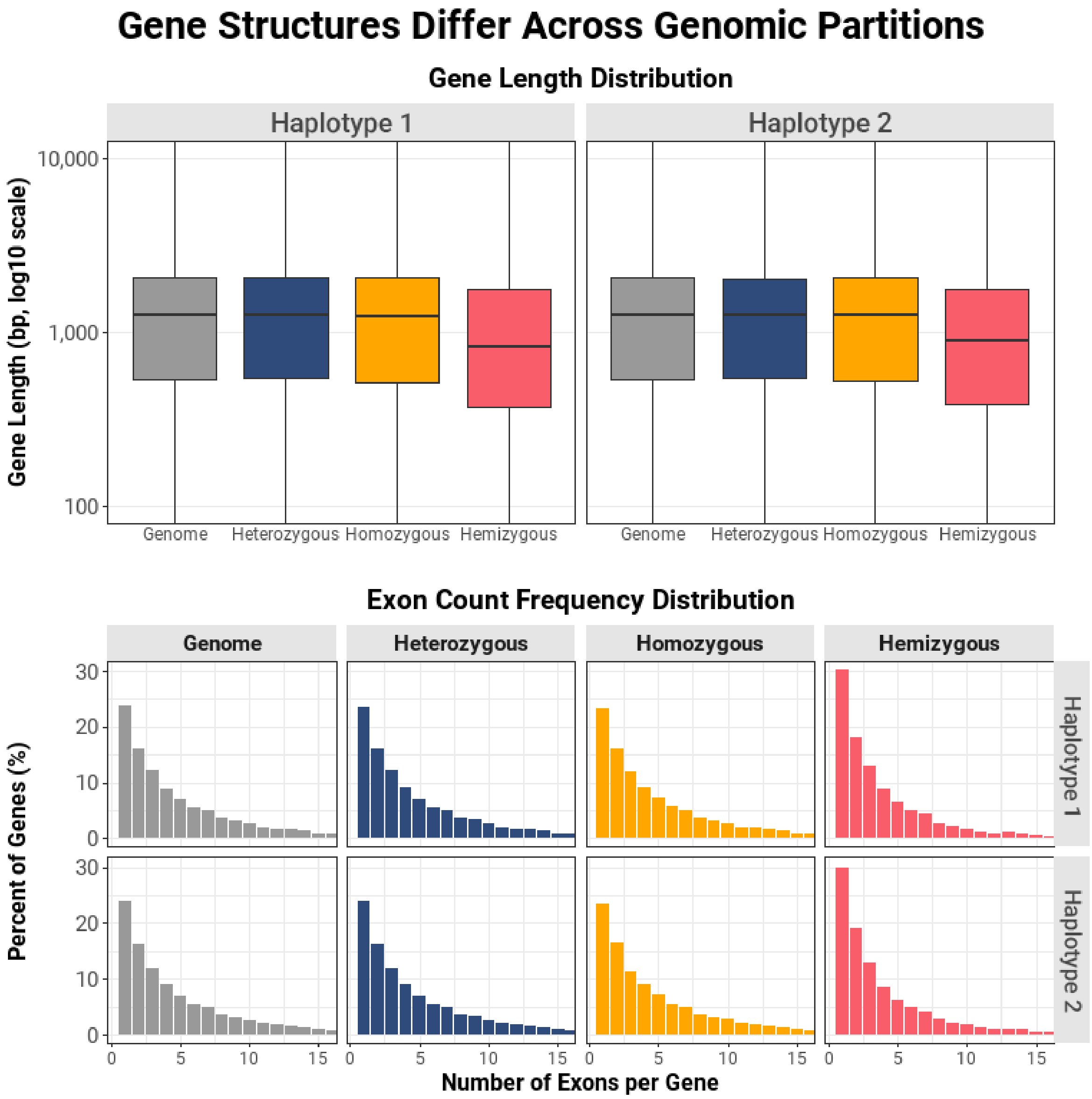


#### Figure S5. Gene Structures Differ Across Genomic Partitions

**(Top panel)** Boxplots showing the distribution of gene lengths (bp, log10 scale) for all genes within each genomic partition, shown separately for each haplotype. **(Bottom panel)** Histograms showing the frequency distribution of the number of exons per gene for each of the four genomic partitions, faceted by haplotype.

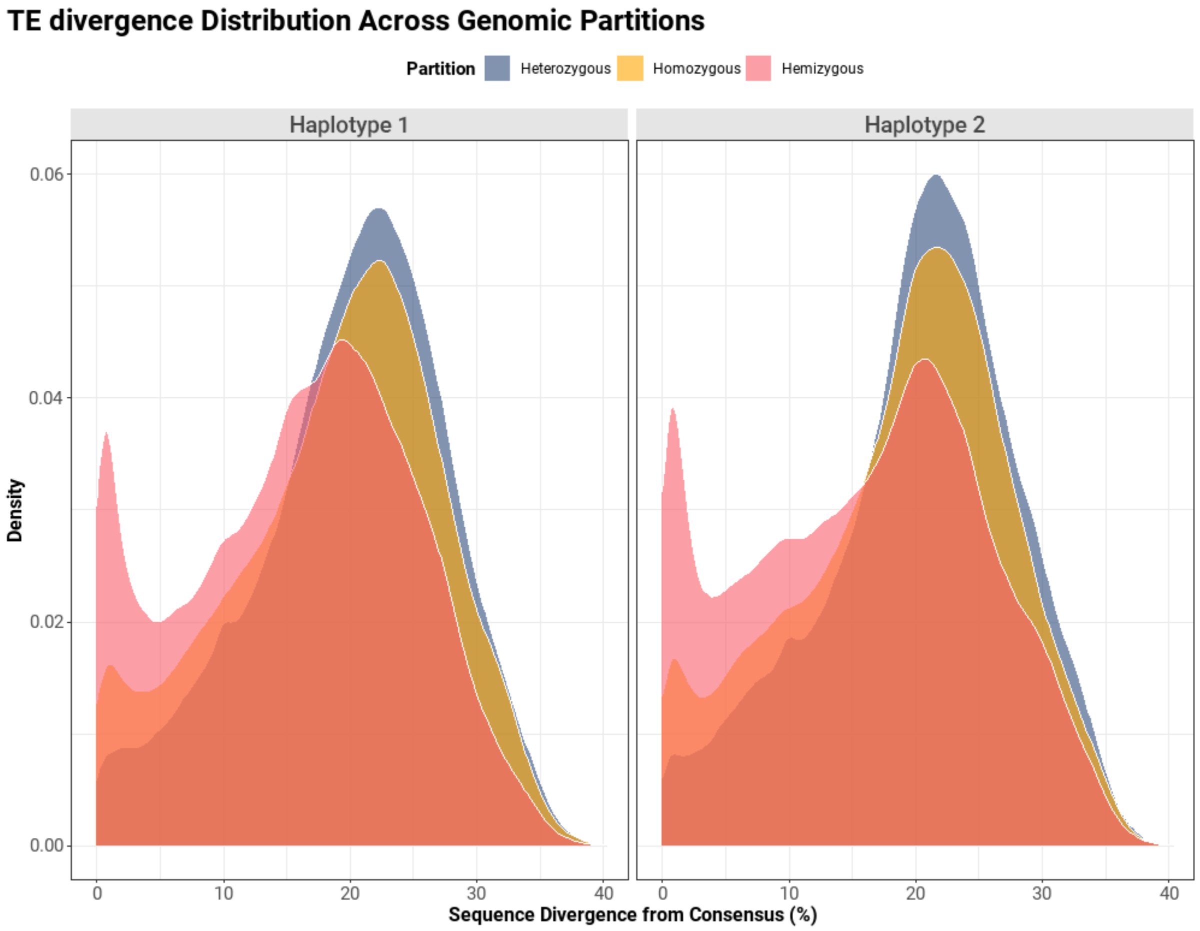


**Figure S6. TE Divergence Distribution Across Genomic Partitions.**

Density plots showing the sequence divergence profiles of transposable elements located within each of the three structural partitions (Heterozygous, Homozygous, Hemizygous). The profiles are shown separately for each haplotype. The sharp peak at low divergence (<5%) for TEs in hemizygous regions is indicative of a recent burst of transposition.

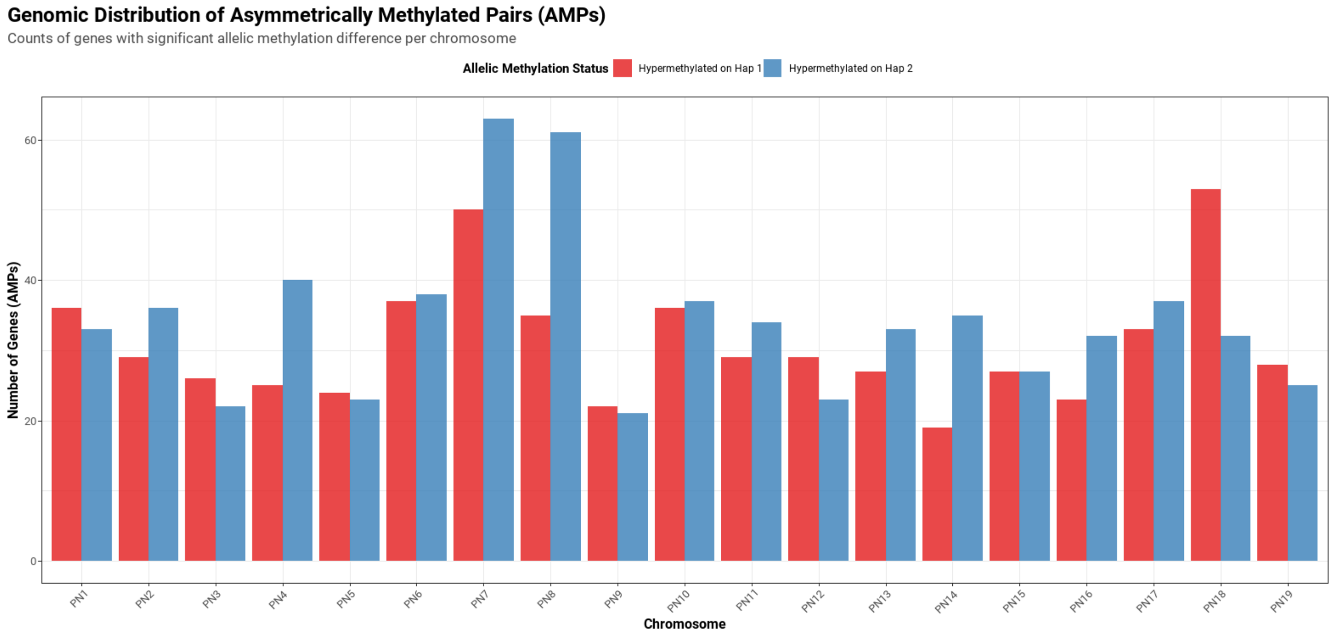


### Figure S7. Genomic Distribution of Asymmetrically Methylated Pairs (AMPs)

Bar chart showing the number of genes classified as AMPs on each of the 19 pseudo-chromosomes. Red bars indicate the count of genes where Haplotype 1 is hypermethylated relative to Haplotype 2. Blue bars indicate the count of genes where Haplotype 2 is hypermethylated relative to Haplotype 1.

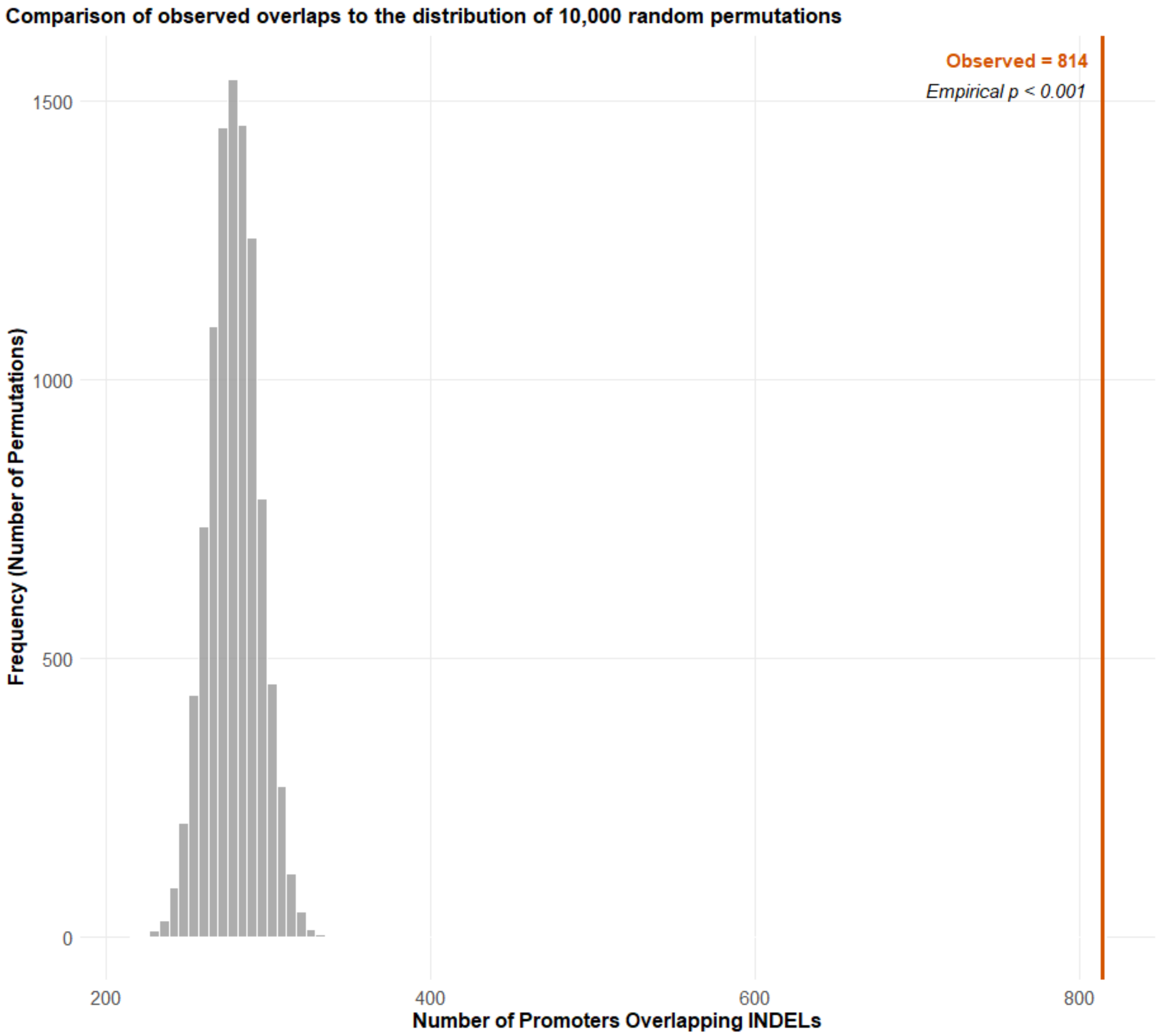


**Figure S8. Permutation Test for Enrichment of AMPs within INDELs.**

Histogram showing the null distribution of the number of overlaps between 10,000 random sets of promoters and large (>100bp) insertion/deletion variants. The red vertical line indicates the observed number of overlaps (814) between the actual set of AMP promoters and the INDELs. The empirical p-value indicates that the observed overlap is significantly greater than expected by chance.

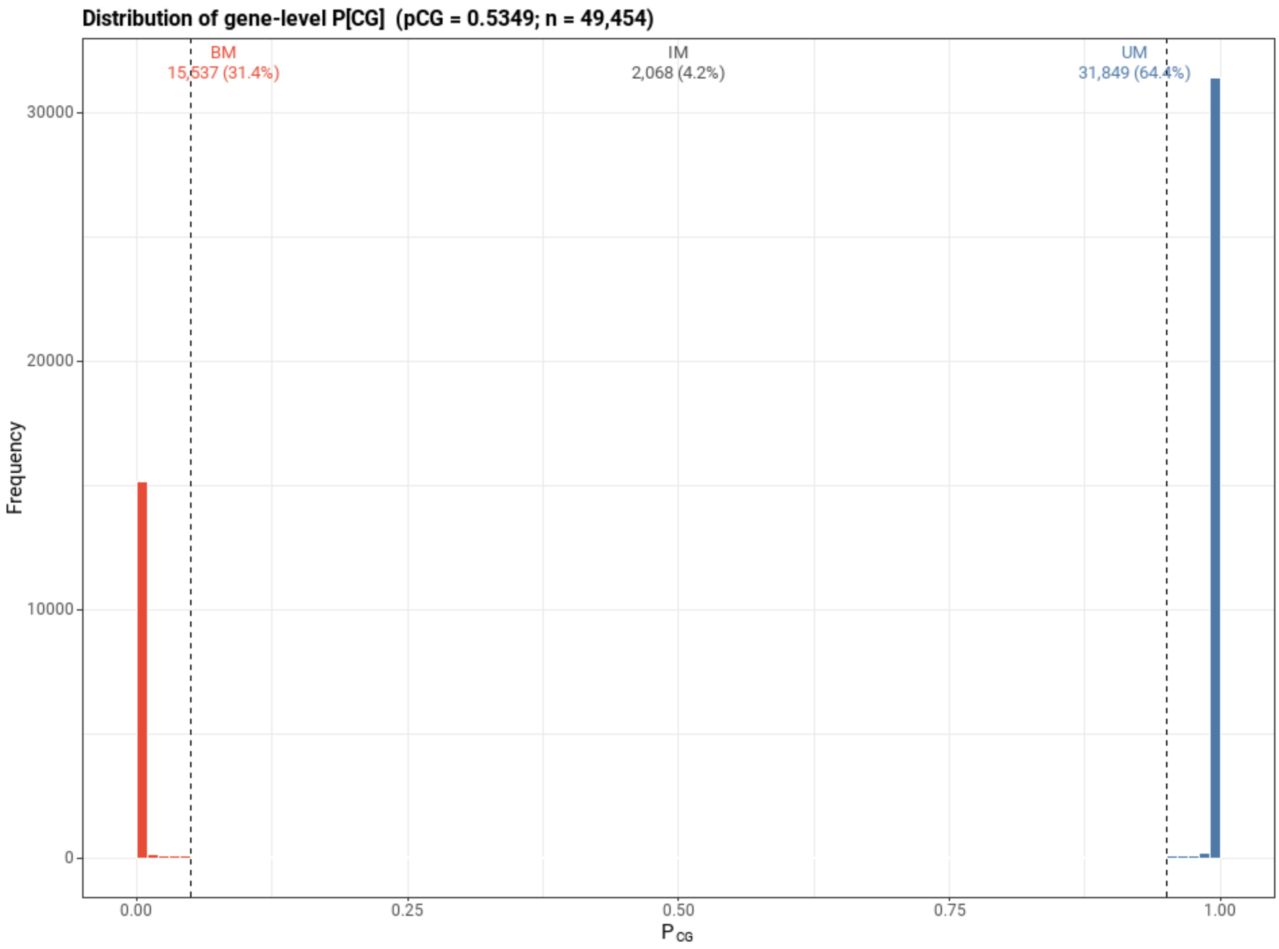


#### Figure S9. Gene Body Methylation (gbM) Classification for the Reference Clone

Histogram showing the distribution of gene-level binomial test P-values for CG methylation ($P_{CG}$) across all 49,454 informative genes (nCG ≥ 20) in the reference clone '20-13 Gm'. The binomial test was used to determine if the number of methylated cytosines observed within a gene is significantly greater than expected by chance, given a background genic CG methylation rate (pCG) of 0.5349.

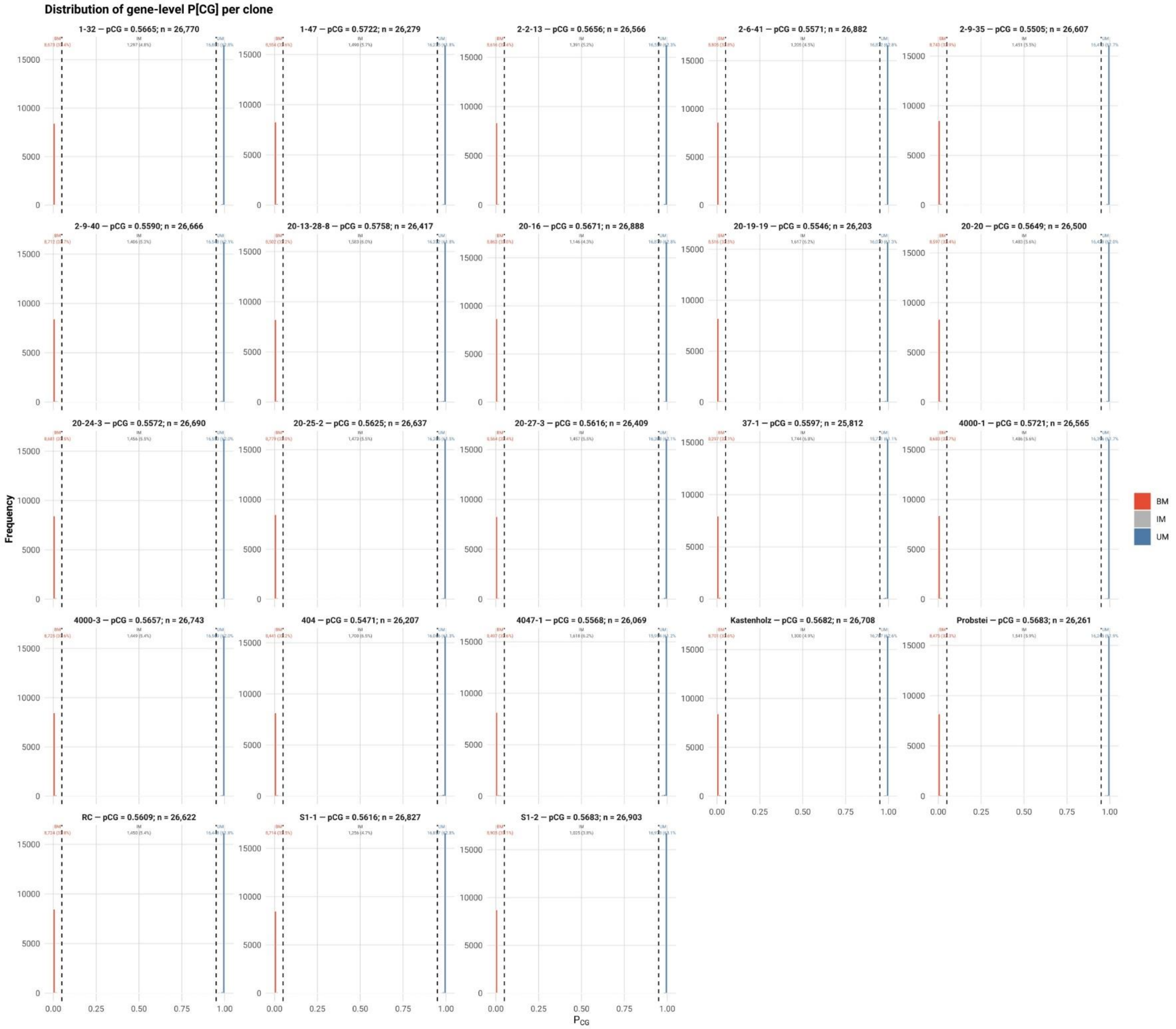


### Figure S10. Per-Clone Gene Body Methylation (gbM) Classification

Faceted plot showing the distribution of gene-level binomial test P-values for CG methylation ($P_{CG}$) for each of the 23 clones individually. For each clone, the binomial test was used to classify informative genes (nCG ≥ 15) as body-methylated (BM; $P_{CG} < 0.05$), intermediately methylated (IM; $0.05 \leq P_{CG} \leq 0.95$), or unmethylated (UM; $P_{CG} > 0.95$). The title of each facet indicates the clone name, its calculated background genic CG methylation rate (pCG), and the total number of informative genes (n) for that clone.

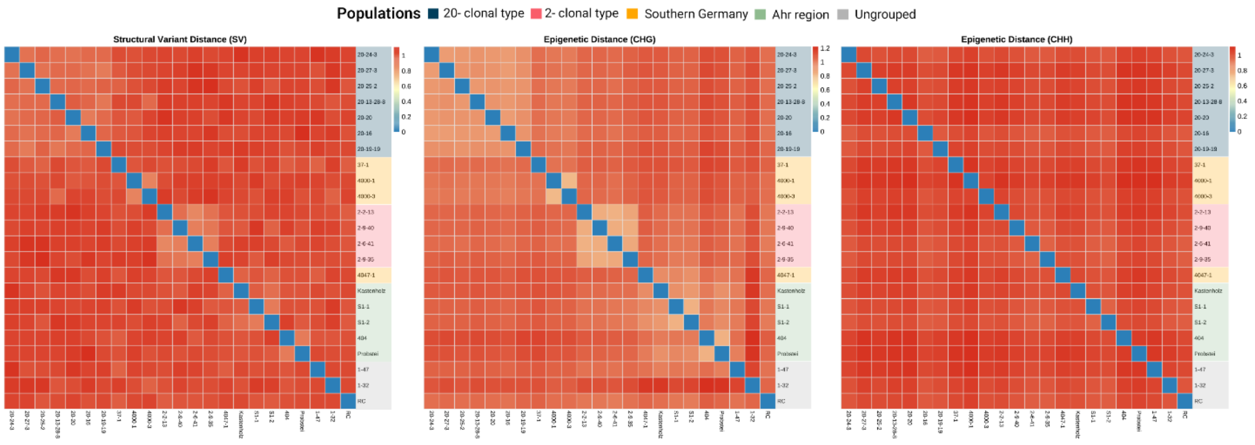


#### Figure S11. Comparison of Distance Metrics across Molecular Layers

Pairwise distance heatmaps (*D=1−Haploid GRM*) for three alternative molecular layers: (Left) Structural Variants (SV), (Middle) CHG-context Differentially Methylated Cytosines, and (Right) CHH-context Differentially Methylated Cytosines. To allow direct comparison with the genetic structure, all three heatmaps are sorted according to the SNP-based phylogenetic order (same as Figure 6).

### Supplementary Tables

#### Table S1. Sequencing statistics for the reference clone '20-13 Gm'.

Metrics for the raw long-read data used for de novo assembly, including read counts, total bases, and N50 read lengths for both PacBio HiFi and Oxford Nanopore Ultra-Long libraries.

#### Table S2. Assembly statistics for the phased pseudo-haplotypes.

Quality metrics for the final scaffolded assembly of Haplotype 1 (PN_1) and Haplotype 2 (PN_2). Columns include total assembly size, contiguity (N50), gene-space completeness (BUSCO scores), and structural accuracy indices (CRAQ S-AQI/R-AQI).

#### Table S3. Gene annotation statistics.

Summary of the functional annotation for both haplotypes. Includes the count of protein-coding genes, exon density, and the proportion of the genome covered by genic sequences.

#### Table S4. Transposable Element (TE) composition.

Detailed quantification of the repetitive fraction of the genome for both haplotypes. The table lists the total base pairs and genomic percentage occupied by major TE superfamilies (LTR, LINE, DNA transposons) and specific subclasses (e.g., Gypsy, Copia).

#### Table S5. Summary of genetic variation between the two haplotypes.

Quantification of the heterozygosity and structural divergence between the two pseudo-haplotypes (PN_1 vs. PN_2) derived from SyRI. This includes the count and total length of SNPs, Insertions, Deletions, and complex rearrangements (Inversions, Translocations, Duplications) defining the diploid landscape.

#### Table S6. Genomic partitioning statistics.

Summary of the three structural partitions defined in the diploid genome. Provides the total size (Mb) and percentage of the genome classified as Heterozygous, Homozygous (RoH), and Functionally Hemizygous for each haplotype.

#### Table S7. Nanopore re-sequencing statistics for the clonal panel.

Sequencing and mapping metrics for the 23 Pinot noir clones. Columns include total read count, mapped read count, mapping efficiency (%), average diploid coverage (X), and mean mapped read length.

#### Table S8. Functional annotation of somatic SNPs.

Summary of the predicted functional impact of the 948 somatic SNPs identified within exonic regions, classified by SnpEff (e.g., Missense, Synonymous, Stop Gained).

### Supplementary bibliography

Liu, Zhongjie; Wang, Nan; Su, Ying; Long, Qiming; Peng, Yanling; Shangguan, Lingfei et al. (2024): Grapevine pangenome facilitates trait genetics and genomic breeding. *Nature Genetics* 56 (12), pp. 2804–2814. DOI: 10.1038/s41588-024-01967-5.

Goel, Manish; Schneeberger, Korbinian (2022): plotsr: visualizing structural similarities and rearrangements between multiple genomes. *Bioinformatics* 38 (10), pp. 2922–2926. DOI: 10.1093/bioinformatics/btac196.

Takuno, Shohei; Seymour, Danelle K.; Gaut, Brandon S. (2017): The Evolutionary Dynamics of Orthologs That Shift in Gene Body Methylation between Arabidopsis Species. *Mol Biol Evol* 34 (6), pp. 1479–1491. DOI: 10.1093/molbev/msx099.
